# Saturated fatty acid chain length shapes B cell lymphoma metabolism and progression

**DOI:** 10.64898/2026.09.18.752627

**Authors:** Audrey L Collin, Marie Pyun, Léa Bourguignon, Bongwe Ngwenyama, Mylène Laflamme, Cassidy Giambagno, Elpida Bokou Gianneli, Isabella Giacomini, Abba Malina, Eric C. Cheung, Fabio Zani, Morgan Craig, Julianna Blagih

## Abstract

B-cell lymphomas reprogram cellular metabolism to sustain their rapid growth and proliferation. Despite their close proximity to adipose tissue and the paradoxical association between obesity and improved lymphoma outcomes, how fatty-acid availability and dietary fat composition influence B-cell lymphoma remains poorly understood. Importantly, saturated fatty acids of different chain lengths have distinct routes of cellular uptake and metabolic fates and may therefore exert divergent effects on tumour growth. Here, we investigated how medium-chain and long-chain saturated fatty acids, their dietary availability, and their transfer from adipocytes shape B-cell lymphoma metabolism and progression. We found that B-cell lymphoma cells were dependent on fatty-acid uptake, activation, and utilization for proliferation. Additionally, we identified adipocytes as microenvironmental players that transfer fatty acids and suppress cell growth. Interestingly, in most B-cell lymphomas, exogenous palmitate limits lymphoma cell proliferation, while MCFA caprylic acid showed only mild effects. Using a syngeneic B-cell lymphoma orthotopic model, we found that an MCFA-rich diet increased lymphoma burden and accelerated lymph-node progression, whereas LCFA-rich and obesogenic diets restricted lymphoma dissemination. Dietary fatty-acid composition also altered T-cell exhaustion and profoundly remodelled the circulating lipidome, with LCFA-rich and obesogenic diets producing highly concordant lipidomic signatures distinct from MCFA feeding. Across B-cell lymphoma cell lines, low ACSL1 expression was associated with increased palmitate sensitivity, which we computationally modelled as a targetable vulnerability in B-cell lymphoma treatment. Taken together, our results show that lipid metabolism is a metabolic vulnerability that can be targeted through dietary intervention.

## Introduction

B-cell lymphomas are highly proliferative malignancies that depend on metabolic adaptation to sustain their energetic and biosynthetic demands[1]. While much of our understanding of lymphoma metabolism has focused on glucose and amino acid utilization, lipids represent an abundant yet comparatively understudied nutrient source within the lymphoma microenvironment[2]. Fatty acids are versatile metabolites, serving not only as energy substrates but also as building blocks for membrane synthesis, lipid storage, and signalling molecules[3]. Aggressive B-cell lymphomas with high CD37 expression depend on fatty acid transport through CD36 and the activation of long-chain fatty acids (LCFAs)[4]. Similarly, a subtype of diffuse large B-cell lymphoma (DLBCL) depends on mitochondrial fatty acid oxidation for proliferation and survival[5]. However, the relevance of lipid metabolism in B-cell lymphomas remains poorly understood.

Within the tumour microenvironment, adipocytes represent a physiologically relevant source of exogenous fatty acids. Adipocytes can release and directly transfer lipids to neighbouring cancer cells, thereby supporting or modifying tumour metabolism[6; 7]. Such interactions have been implicated in several solid and haematological malignancies[6-8], yet metabolic communication between adipocytes and B-cell lymphoma remains poorly defined. This relationship is particularly intriguing given epidemiological observations linking higher body mass index to improved treatment responses or survival in some B-cell lymphomas, despite obesity being associated with higher incidence and poorer outcomes in many other cancers[9]. These apparently paradoxical observations raise the possibility that the lipid-rich environment associated with increased adiposity may have anti-tumorigenic effects on lymphoma cells.

Diet represents another major determinant of the extracellular lipid environment[10]. Dietary fat composition influences circulating fatty acids and complex lipid species and can thereby alter the metabolic substrates available to both tumour and immune cells. However, most studies investigating diet and blood cancers have focused on total dietary fat, obesity, or caloric excess, making it difficult to distinguish the consequences of increased adiposity from those of specific dietary fatty acids[11-16]. In particular, whether the chain length of dietary saturated fatty acids influences lymphoma growth and dissemination remains unknown. Understanding this distinction may also help reconcile the complex relationship between adiposity and clinical outcomes in B-cell lymphoma.

Here, we investigated how exogenous saturated fatty acids influence B-cell lymphoma metabolism and progression. We first defined lymphoma dependencies on pathways controlling fatty-acid uptake, activation, and mitochondrial utilization and determined whether sensitivity to MCFAs and LCFAs was associated with mitochondrial metabolic fitness. We next examined adipocytes as a physiologically relevant source of fatty acids and assessed metabolic communication between adipocytes and lymphoma cells. Finally, using isocaloric diets differing in saturated fatty acid chain length, we determined how dietary MCFAs and LCFAs remodel the circulating lipidome, lymphoma progression, and the tumour immune microenvironment. Finally, we used a mathematical model to predict the effects of an LCFA-enriched diet on B-cell lymphoma growth, showing that this dietary modification could provide potential survival benefits for certain B-cell lymphoma subtypes. Together, Our findings reveal that saturated fatty acid chain length is a critical determinant of lymphoma response to the extracellular lipid environment and identify lipid-handling capacity as a potential vulnerability that could inform metabolically guided nutritional strategies in B-cell lymphoma.

## Materials and Methods

### Animals

Mice were housed under specific pathogen-free (SPF) conditions at the Centre de Recherche de l’Hôpital Maisonneuve Rosemont (CR-HMR) in accordance with institutional and national guidelines for the care and use of laboratory animals. Animals were maintained under controlled environmental conditions with a 12-h light/12-h dark cycle, controlled temperature and humidity, and ad libitum access to water and the indicated experimental diets. Mice were housed in individually ventilated cages with appropriate bedding and environmental enrichment. We routinely monitored animals for health and welfare throughout the experimental period. Unless otherwise indicated, age- and sex-matched mice were randomly assigned to experimental groups. Experimental diets were initiated at 6 weeks and were provided ad libitum for the indicated duration. The following diets were purchased from Inotive: Purified diet (TD.97184), 3% CocoaB, 3% Palm Oil diet [93G, VI, R] (TD.250159), 6% CCO Diet [93G, VI, B] (TD.250160), and Adjusted Calories Diet (TD.06414). Body weight and general health were monitored throughout dietary interventions. Where applicable, humane endpoints were predefined according to the approved animal-use protocol. All animal procedures were reviewed and approved by the Institutional Animal Care and Use Committee of CR-HMR (protocol no. 2025-3841 and 2025-4039) and were performed in accordance with the guidelines of the Canadian Council on Animal Care (CCAC).

### Cell Culture

#### B cell lymphoma cell lines

Six human Burkitt’s lymphoma cell lines were used, including three EBV⁺ cell lines (Daudi, Namalwa, Raji) and three EBV⁻ cell lines (BJAB Cas9 E3, DG-75, Ramos). The cell lines were maintained in complete RPMI-1640 medium (Wisent Multicell), supplemented with 10% fetal bovine serum (FBS; Wisent Multicell), 1% penicillin-streptomycin (P/S; 10,000 U/mL; Wisent Multicell), and 0.1% 2-mercaptoethanol (55 mM; Gibco). Subcultured cultures every 48 to 72 hours at a 1:5 ratio.

The murine 2F5 cell line (B lymphocytes) was cultured on a layer of irradiated murine embryonic fibroblasts in a medium composed of 50% DMEM and 50% complete IMDM (Wisent Multicell), supplemented with 10% FBS and 1% P/S. We subcultured cultures every 48 to 72 hours at a 1:5 ratio. cells were incubated at 37°C in an incubator humidified with 5% CO₂.

#### Maintenance and differentiation of 3T3-L1 pre-adipocytes

3T3-L1 preadipocytes were maintained DMEM supplemented with 10% FBS and 1% P/S at 37°C in a humidified atmosphere containing 5% CO₂. Cells were routinely passaged before reaching complete confluence and were maintained at subconfluent densities to preserve their differentiation capacity. The culture medium was replaced every 2–3 days. For differentiation, 3T3-L1 cells were seeded in 24-well plates and cultured to ∼80% confluence. Adipocyte differentiation was induced with complete DMEM supplemented with 0.5 mM IBMX (Acros Organics), 0.5 µM dexamethasone (Gibco), 10 µg/mL insulin (Sigma-Aldrich), and 20 µM troglitazone (Sigma).On days 4, 6, and 8, the medium was replaced with complete DMEM supplemented with 10 µg/mL insulin. On day 10, the medium was replaced with standard complete DMEM. The 3T3-L1 cell line (murine pre-adipocytes) was maintained in complete DMEM (Wisent Multicell) supplemented with 10% FBS and 1% P/S. cells were incubated at 37°C in an incubator humidified with 5% CO₂.

#### Isolation and differentiation of white pre-adipocytes

White adipose tissue (WAT) was collected from visceral and subcutaneous adipose deposits in C57BL/6 mice and placed in DMEM/F12 supplemented with 1% P/S. WAT was weighed, minced into small pieces, and digested in HBSS containing 1% FBS, penicillin/streptomycin, and 0.15% (w/v) collagenase D at 6 mL of digestion buffer per gram of tissue. Digestion was performed for 30 min at 37°C with orbital shaking. The digested tissue was filtered through a 300-µm cell strainer and washed with PBS, then centrifuged at 400 × g for 10 min at room temperature. The resulting pellet was treated with 1× RBC lysis buffer for 2 minutes at room temperature, followed by addition of DMEM/F12 containing 10% FBS and penicillin/streptomycin, then centrifuged at 400 × g for 10 min. Cells were resuspended in culture medium and filtered through a 40-µm cell strainer before counting using trypan blue exclusion. Cells were seeded at 3 × 10^5^ cells/well in 6-well plates in DMEM/F12 supplemented with 10% FBS and penicillin/streptomycin (2 mL/well) and maintained at 37°C in 5% CO₂. Culture medium was replaced every 2 days, and cells were allowed to reach confluence over approximately 7 days before differentiation or subsequent experiments following the protocol for 3T3-L.

### B cell lymphoma proliferation assays

Proliferation assays were performed in 96-well U-bottom plates, in triplicate. Cells were seeded at the following densities: 50,000 cells/well (BJAB Cas9 E3, Daudi, DG-75, Raji, 2F5), 80,000 cells/well (Ramos), and 100,000 cells/well (Namalwa). Cells for counting on days 3 and 6 were seeded in separate wells. On day 3, cells for day 6 counting were transferred to a 48-well plate, and the medium volume was adjusted. We counted cells by trypan blue exclusion (1:1) using a DeNovix cell counter. Unless otherwise indicated, we took measurements on days 3 and 6. To assess the effects of saturated fatty-acid availability on cell proliferation, cells were cultured in standard growth medium or medium supplemented with BSA-conjugated caprylic acid (C8; Cayman Chemical) or BSA-conjugated palmitic acid (C16; Cayman Chemical) at final concentrations of 50 or 100 µM. Cells were cultured for up to 6 days, with viable cell numbers determined as described above. For experiments assessing the contribution of fatty-acid uptake or metabolism, cells were cultured under two conditions: standard growth medium and in combination with the indicated inhibitors. Cells were cultured for up to 6 days and viable cell numbers were determined at the indicated time points. The figures specify the inhibitors and their concentrations for each experiment.

#### Proliferation with adipocytes or adipocyte conditioned media (ACM)

Differentiated 3T3-L1 adipocytes (24-well plates) (see section above) were co-cultured with Burkitt’s Lymphoma cells in triplicate at the following densities: 100,000 cells/well (BJAB Cas9 E3, Daudi, DG-75, Raji), 160,000 cells/well (Ramos), and 200,000 cells/well (Namalwa or EμMyc cells). BL cells were co-cultured with the 3T3-L1 cells for 72 hours. Cell counts were performed every 24 hours.

Adipocyte-conditioned medium (ACM) was collected from fully differentiated 3T3-L1 adipocytes. Following confirmation of adipocyte differentiation, the existing culture medium was removed and replaced with fresh culture medium. Cells were conditioned for 24 h under standard culture conditions (37°C, 5% CO₂), after which the medium was collected and used as ACM for subsequent experiments. Unless otherwise indicated, ACM was mixed at a 1:1 ratio with complete RPMI-1640 medium for treatment of B-cell lymphoma cells.

### Cellular bioenergetics

#### MitoStress Test

Cells were pre-incubated for 24 hours at 4,000,000 cells/well in 6-well plates in complete RPMI-1640. The Seahorse XF assay medium was prepared from Seahorse XF Base Medium supplemented with 2 mM sodium pyruvate, 2 mM glutamine, and 25 mM glucose (Agilent Seahorse). Mitochondrial modulators were prepared to a final concentration of 1 µM: oligomycin (ATP synthase inhibitor; Cayman Chem), FCCP (uncoupler), and antimycin A (complex III inhibitor). A cell suspension was seeded onto a 96-well Seahorse XF plate pre-coated with poly-D-lysine at a density of 2,000,000 cells/mL. The cells were centrifuged and then incubated for 1 hour at 37°C without CO₂. The modulators were loaded into the injection ports of the sensor cartridge. Calibration and data acquisition were performed using an Agilent Seahorse XF analyzer, and the data were then processed with Agilent Seahorse XFp v2.5.3 software.

#### Palmitate Oxidation Test

On day 1, the sensor cartridges according to the manufacturer’s (Agilent) recommendations. The cells were incubated overnight at 2,000,000 cells/well (6-well plates) in a substrate-limited growth medium containing 0.5 mM glucose, 1.0 mM glutamine, 1% FBS, and 0.5 mM L-carnitine (Sigma-Aldrich), at 37 °C and 5% CO₂. On day 2, the water in the sensor cartridge was replaced with the XF calibrating solution, then incubated at 37°C without CO₂ for 1 hour. The cells were counted and seeded in 96-well Seahorse XF plates (pre-coated with poly-D-lysine) at 100,000 cells/well in substrate-limited assay medium containing 2.0 mM glucose and 0.5 mM L-carnitine, then incubated at 37°C without CO₂ for 1 hour. The mitochondrial modulators (oligomycin, FCCP, and antimycin A) were prepared to a final concentration of 2.5 µM per well and loaded into the cartridge before calibration. After calibration, we added BSA palmitate to a final concentration of 50 µM to the cells immediately before the assay began.

### Flow Cytometry

#### Viability staining

The cells were incubated for 10 minutes at 4°C with 50 µL of eFluor 780 viability dye diluted 1:1000 in 1X PBS. The cells were then washed with 1X PBS and centrifuged at 1300 rpm for 5 minutes.

#### Mitochondrial mass and potential

Cells were seeded at 500,000 cells/mL (BJAB Cas9 E3, Daudi, DG-75, Raji), 800,000 cells/mL (Ramos), or 1,000,000 cells/mL (Namalwa) and incubated for 24 hours in their growth medium. Cells were harvested by centrifugation (1300 rpm, 5 minutes), and viability staining was done as described above. The cells were then incubated for 30 minutes at 37°C in the dark with MitoTracker Green (20 nM) and MitoTracker Deep Red (50 nM) (Invitrogen), diluted in 1X PBS. After washing, the cells were resuspended in their growth medium and stored at 4°C until acquisition.

#### Absorption of saturated fatty acids and lipid droplets

Cells were seeded at 500,000 cells/mL (BJAB Cas9 E3, Daudi, DG-75, Raji), 800,000 cells/mL (Ramos), or 1,000,000 cells/mL (Namalwa) and incubated for 48 hours. On day 3, the medium was replaced with fresh medium containing either BODIPY 558/568 C12 (2 µM; Invitrogen) or BODIPY FL C16 (1 µM; Invitrogen), and the cells were incubated for an additional 24 hours. Cells were seeded at the densities described above and incubated for 24 hours in standard medium, followed by staining with the lipid droplet marker BODIPY 4993/503 (0.5 µM), followed by a 2-hour incubation at 37°C and 5% CO_2_.

#### Mitochondrial ROS and cellular ROS

Cells were seeded at the densities described above and incubated for 72 hours in standard medium, medium enriched with BSA-caprylic acid (50 µM), or medium enriched with BSA-palmitate (50 µM). After harvesting (1300 rpm, 5 minutes), we incubated the cells for 30 minutes at 37°C in the dark with MitoSOX (1 µM) or CellROX (1 µM; Invitrogen).

#### Transfer of fatty acids from adipocytes to lymphoma cells

Fatty-acid transfer from adipocytes to lymphoma cells was assessed using BODIPY 558/568 C12 and BODIPY FL C16 as fluorescent fatty-acid analogues. Fully differentiated adipocytes were incubated overnight with BODIPY 558/568 C12 (2 µM) and/or BODIPY FL C16 (1 µM), washed with 1× PBS to remove unincorporated probe, and subsequently co-cultured with lymphoma cells. For experiments using 3T3-L1 adipocytes, human B-cell lymphoma cells were added at 5 × 10⁵ cells/well for BJAB Cas9 E3, Daudi, DG-75, and Raji; 8 × 10⁵ cells/well for Ramos; and 1 × 10⁶ cells/well for Namalwa. For primary adipocyte experiments, adipocytes differentiated from subcutaneous (SAT) or visceral adipose tissue (VAT)-derived preadipocytes were co-cultured with Eμ-Myc cells (5 × 10⁵ cells/well) in 24-well plates for 24 h. Following co-culture, lymphoma cells were collected by centrifugation (1,300 rpm, 5 min), washed, and analyzed by flow cytometry to quantify adipocyte-to-lymphoma cell fatty-acid transfer.

### Western Blot

The cells were seeded at 2x10^6^ cells/well (6-well plates) and incubated for 24 hours. Protein lysates were prepared in RIPA buffer (Millipore Sigma) supplemented with a protease/phosphatase inhibitor cocktail (Cell Signaling; 1:100), 10% SDS (1:100), and 1 M benzamidine hydrochloride (1:2,000; Millipore Sigma). The lysates were vortexed and then mechanically sheared using an insulin syringe (BD). After centrifugation, we determined protein concentration by Bradford assay. For the analysis, 15 µg of protein per sample was separated on a 10% SDS-PAGE gel and then transferred to PVDF membranes by wet transfer. After blocking, the membranes were incubated with primary antibodies against fatty acid and lipid metabolism proteins, as well as an anti-β-actin antibody as a loading control. After incubation with HRP-conjugated secondary antibodies, the signals were detected by chemiluminescence using the West Femto substrate and acquired on an Azure c6000 imaging system.

### EμMyc orthotopic model

Male C57BL/6J mice were purchased at 6 weeks of age and allowed to acclimatise for 1 week (Jackson Laboratory). At 6 weeks of age, mice were randomly assigned to one of four experimental diets: (i) a control diet containing 17.4% fat; (ii) a medium-chain fatty acid (MCFA)-enriched diet, in which 6% of the control diet was replaced with coconut oil; (iii) a long-chain fatty acid (LCFA)-enriched diet, in which 3% of the control diet was replaced with cocoa butter and 3% with palm oil; or (iv) a high-fat obesogenic diet. Mice were maintained on their respective diets throughout the experiment, and body weight was recorded weekly. After 28 days of dietary intervention, mice were intravenously injected with 1 × 10⁶ Eμ-Myc lymphoma cells. Mice remained on their respective diets for an additional 12 days and were then euthanized for endpoint analyses. Lymphoma burden and dissemination were assessed by spleen weight and by quantification and measurement of enlarged lymph nodes at endpoint.

### Tumour immune phenotyping: Splenic immune phenotyping

At experimental endpoint, spleens were collected and processed into single-cell suspensions for flow cytometric analysis of immune-cell populations. Spleens were mechanically dissociated through a 70-µm cell strainer and washed with PBS containing 2% FBS. Red blood cells were lysed using ammonium-chloride-potassium (ACK) lysis buffer, after which cells were washed, resuspended in flow cytometry staining buffer, and counted.

For surface immunophenotyping, cells were incubated with Fc receptor-blocking antibody followed by fluorophore-conjugated antibodies against the indicated immune-cell surface markers. eBioscience™ Fixable Viability Dye eFluor™ 780 (1/1000 dilution) was used to exclude dead cells from subsequent analyses. Surface staining was performed in PBS, and 1% BSA; all antibodies were used at a dilution of 1/300. Samples were acquired on a BD LSRFortessa X-20 flow cytometer (BD Biosciences) and analyzed using FlowJo software (BD Biosciences). We identified immune-cell populations from singlet, viable leukocytes using the gating strategies shown in the corresponding figures.

### In vitro CD8 T cell exhaustion assay

Splenocytes were collected from the spleen of C57BL/6 mice and suspended as single-cell suspensions. CD8^+^ or CD4 ^+^ T Cells were isolated using their respective negative selection kits from Stem Cell according to the manufacture’s protocol. Primary T cells were cultured for 6 days in T-cell medium (IMDM supplemented with 10% FBS, 1% P/S, and 50µM β-mercaptoethanol). On day 0, culture plates were coated with 5 µg/mL of anti-CD3 (Biolegend, 145-2C11, 1015436) and 2 µg/mL anti-CD28 (Biolegend, 37.51, 14-0281-85) overnight at 4°C or for 2 h at 37°C. The isolated T cells were then seeded at 1 × 10^6^ cells/well in the coated plates with 10 ng/mL IL-2 and cultured for 3 days. On day 3, cells were harvested, counted, and replated at 1 × 10^6 cells/well in anti-CD3 (5 µg/mL) coated plates and 10 ng/mL IL-2. On day 5, cells were replated with the same conditions as day 3. On day 6, the cells were stimulated with PMA/ionomycin and GolgiStop for 4 h at 37°C and 5% CO₂, followed by intracellular cytokine staining for IFNγ and TNFα.

### Serum lipidomic profiling

Serum samples collected from mice following dietary treatment were submitted to Linearis Labs (Canada) for targeted metabolomic and lipidomic profiling using the GigaKit™ Metabolomic Panel. The assay quantified up to 1,780 metabolites across 38 biochemical classes using liquid chromatography-tandem mass spectrometry (LC-MS/MS) and direct flow injection-tandem mass spectrometry (DFI-MS/MS). Briefly, serum samples were subjected to pre-column derivatization using phenylisothiocyanate (PITC) for amine-containing metabolites and 3-nitrophenylhydrazine (3-NPH) for organic acids and selected lipids. Derivatized samples were analyzed in multiple reaction monitoring (MRM) mode using a Waters UPLC Acquity I-Class system coupled to a Xevo TQ-Absolute mass spectrometer, with five analytical runs performed per sample. Data acquisition, processing, and quantification were performed using Waters Connect software. Metabolites analyzed by LC-MS/MS were quantified using seven-point calibration curves based on analyte-to-internal standard (ISTD) peak intensity ratios, whereas lipids and acylcarnitines were semi-quantitatively assessed by DFI-MS using single-point calibration with representative analyte and ISTD standards for each chemical class. Quality control pooled serum samples were included in each analytical batch to monitor measurement variability and facilitate data normalization. The analytical laboratory assessed data quality using an automated quality control pipeline.

### Mathematical model of tumour growth and inhibitory effect of LCFA on Burkitt lymphoma proliferation

Based on observed growth patterns in our in vitro data (Fig. 2A and 2B) and previous murine studies[17; 18], we used the standard logistic growth model[19] to represent BL growth over time. Let *V* denote tumour volume (mm^3^); this model is given by

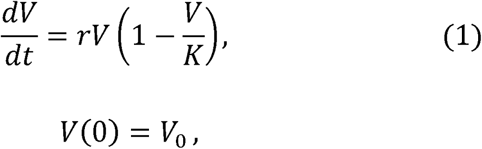

where *r* represents the intrinsic tumour growth rate, *K* denotes the maximum tumour volume (i.e., tumour carrying capacity), and *V*_0_ is the initial tumour size[20]. We used a standard inhibitory dose-response curve[21] to represent the observed inhibitory effects of LCFA on BL growth (Fig. 2B). In this model, LCFAs act as the inhibitor, with response measured as the fold change from untreated tumour growth, i.e.,

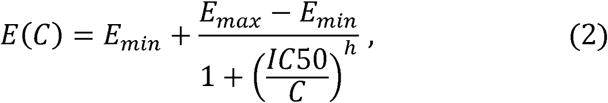

where *E*(*C*) is the effect of LCFAs on cell proliferation at concentration *C*, *E_min_* and *E_max_* are the minimal (i.e., *C* = 0) and maximal (i.e., *C* → ∞) effects, respectively, *IC*50 is the concentration producing 50% of maximal inhibition, and *h* is the Hill coefficient representing the steepness of the curve[22].

All computations in this study were performed in MATLAB version R2025b, Update 3[23].

#### Parameter estimation

We estimated the parameters in Eqs. 1 and 2 by minimizing the residual sum of squares (RSS), i.e.,

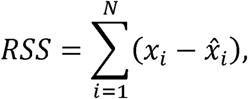

where *N* is the total number of observed data points, *x* denotes the model prediction, and *x̂* represents the observed data. As RSS is sensitive to local minima, we used the in-built *MultiStart* function in Matlab[23] and generated 20 runs with randomly sampled initial parameter values, retaining only the best-fitting solution.

We used data from Huang et al.[17] and Zhang et al.[18], who reported tumour growth in BALB/c nude mice after inoculation with Namalwa (low ACSL1 expression; 5 mice)[17] and Ramos (high ACSL1 expression; 7 mice)[18] BL cells to estimate the intrinsic tumour growth rate, *r*, and the maximal tumour volume, *K*, in Eq. (1). As neither study reported the initial tumour volume, *V*_0_, we first estimated plausible values for each timeseries using a grid search over 25 logarithmically spaced candidates ranging from 10^-6^ mm^3^ (a negligible volume) to a theoretical maximum; the latter was calculated from the number of cells initially inoculated in each mouse, assuming that 10^6^ cells = 1 mm^3^, as in previous studies[24]. Then, for each candidate *V*_0_, *r* and *K* were estimated by minimizing the RSS, and the five candidates yielding the lowest RSS values were retained. Selecting five candidates, rather than one, allowed us to capture additional variability within each dataset.

For each of the retained *V*_0_, we estimated 95% confidence intervals (CIs) for the values of *r* and *K* using the profile likelihood method[25] (see Fig. S8 and Table S2 for results). From the profile likelihood, we calculated the probability density function (pdf) of each parameter within their respective CI through normalization of the likelihood over the confidence interval. The profile likelihood method also doubled as a verification of parameter practical identifiability.

We used the same *MultiStart* procedure and minimization of RSS to estimate the parameters in the inhibitory-response model (Eq. 2). To quantify the effect of LCFAs on BL proliferation, we considered four of the six lines in our in vitro cell culture data (Fig. 2B): Namalwa and DG-75 (low ACSL1 expression), and Daudi and Ramos (high ACSL1 expression). For each LCFA treatment and expression group, we calculated the fold change between mean day-6 cell count between treated and untreated control. Assuming no effect in normal conditions, we fixed *E_min_* to be 1, with *E_max_* constrained between 0 and 1 (corresponding to the range of possible fold change values).

#### Generation of virtual Burkitt Lymphoma patient populations

Using our calibrated mathematical model (Eqs. 1 and 2), we generated two virtual populations corresponding to high and low ACSL1 expression groups. Each virtual population comprised 1,000 individuals, defined to be an (*r*, *K*)-parameter pair (Eq. 1) sampled from the profile likelihood curves generated during parameter estimation (see above). Parameter pairs were obtained by randomly selecting one of the five *V*_0_ values, then sampling r and *K* from their associated CIs, according to the corresponding pdf. Sampling following these pdfs was performed using inverse transform sampling[26].

#### Simulation of increased dietary LCFA intake in virtual populations

To simulate the potential effects of LCFA-enriched diets on Burkitt lymphoma growth, we considered three dietary interventions, differing in their extent of LCFA exposure, in our virtual populations: 1) a standard diet, consisting of LCFA concentrations ≤5 μM to reflect background exposure; 2) Diet 1, a moderate increase in dietary LCFA, with concentrations peaking between 25 and 35 μM, a range chosen to encompass the IC50 of the low-ACSL1-expression group; and 3) Diet 2, a substantial increase in dietary LCFA, with concentrations peaking between 90 and 100 μM. We note that these concentrations are neither excessive nor toxic: circulating LCFA concentrations in mice can safely reach up to ∼100 μM, with some studies finding levels as high as ∼300 μM.[27-29]

For each diet, we simulated three daily meals which evenly spaced across a 12-hour feeding period followed by a 12-hour fast. To approximate realistic in vivo exposure, we modelled LCFAs as fluctuating over time such that their concentration would rise instantaneously after each meal, then decay exponentially at rate *a* between meals. For each individual, the LCFA concentration, *C*, of each meal was drawn from a uniform distribution over the corresponding diet’s range. The exponential decay rate, *a*, was sampled per individual from *Unif[10, 20]*, chosen so that LCFA concentrations returned to zero during the fasting period under Diet 1 and accumulated only modestly across consecutive meals. This pharmacokinetic model is thus described by:

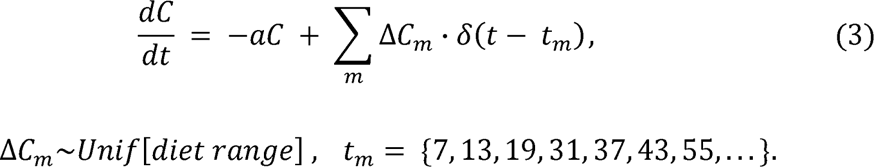

The effects of these time-varying LCFA concentrations were then linked to tumour growth by scaling *r* in Eq. 1 by *E*(*C*) (Equation 2), with *C* fluctuating per Eq. 3:

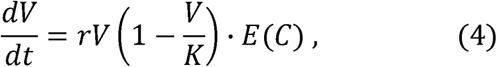

We used the integrated model to simulate tumour growth in each of the 1,000 individuals in both virtual populations under each dietary condition. Tumour growth was simulated over 50 days, a period selected because it was sufficient for all tumours under the standard diet to reach their maximum predicted volume.

### Predicting survival in virtual cohorts

To express predicted tumour growth as a survival probability, we used a lethality threshold of 5200 mm^3^, chosen to so that the resulting Kaplan-Meier survival curves generated under the standard diet would reasonably reproduce Burkitt Lymphoma survival trends. Using this threshold, the low- and high-ACSL1 groups virtual populations reached endpoint survival rates of 52.3% and 95.5%, respectively, compared with approximately 67% and 86% in the PRECOG cohort (Fig. 4C).

## Results

### B-cell lymphoma cells are vulnerable to fatty acid metabolism

We first investigated the general metabolic profile of 6 Burkitt lymphoma cell lines: three with past Epstein-Barr virus (EBV) infection (Daudi, Namalwa and Raji)[30] and three without (BJAB, DG75, and Ramos). We measured energetic status using the Seahorse Mitostress Test, where oxygen consumption rate (OCR) served as a proxy for mitochondrial activity and extracellular acidification rate (ECAR) as a measure of glycolysis. Stratifying the cell lines by OCR and ECAR values revealed no major differences attributable to prior EBV infection, including differences in spare respiratory capacity, a measure of mitochondrial fitness (**Figure 1A and B**). EBV infection in B cells reprograms mitochondria[31]; therefore, we evaluated the mitochondrial parameters: mitochondrial load with MitoTracker Green and membrane potential with MitoTracker Red. Raji (EBV-positive history) was the only cell line exhibiting a distinct mitochondrial profile (**Figure S1A**). Given that EBV can remodel lipid metabolism[32], we assessed lipid droplet formation with the BODIPY 493/503 dye and did not observe EBV-dependent alterations **(Figure 1C**). We also found no correlation between EBV status and CD36 expression (**Figure S1B**).

**Figure 1:**
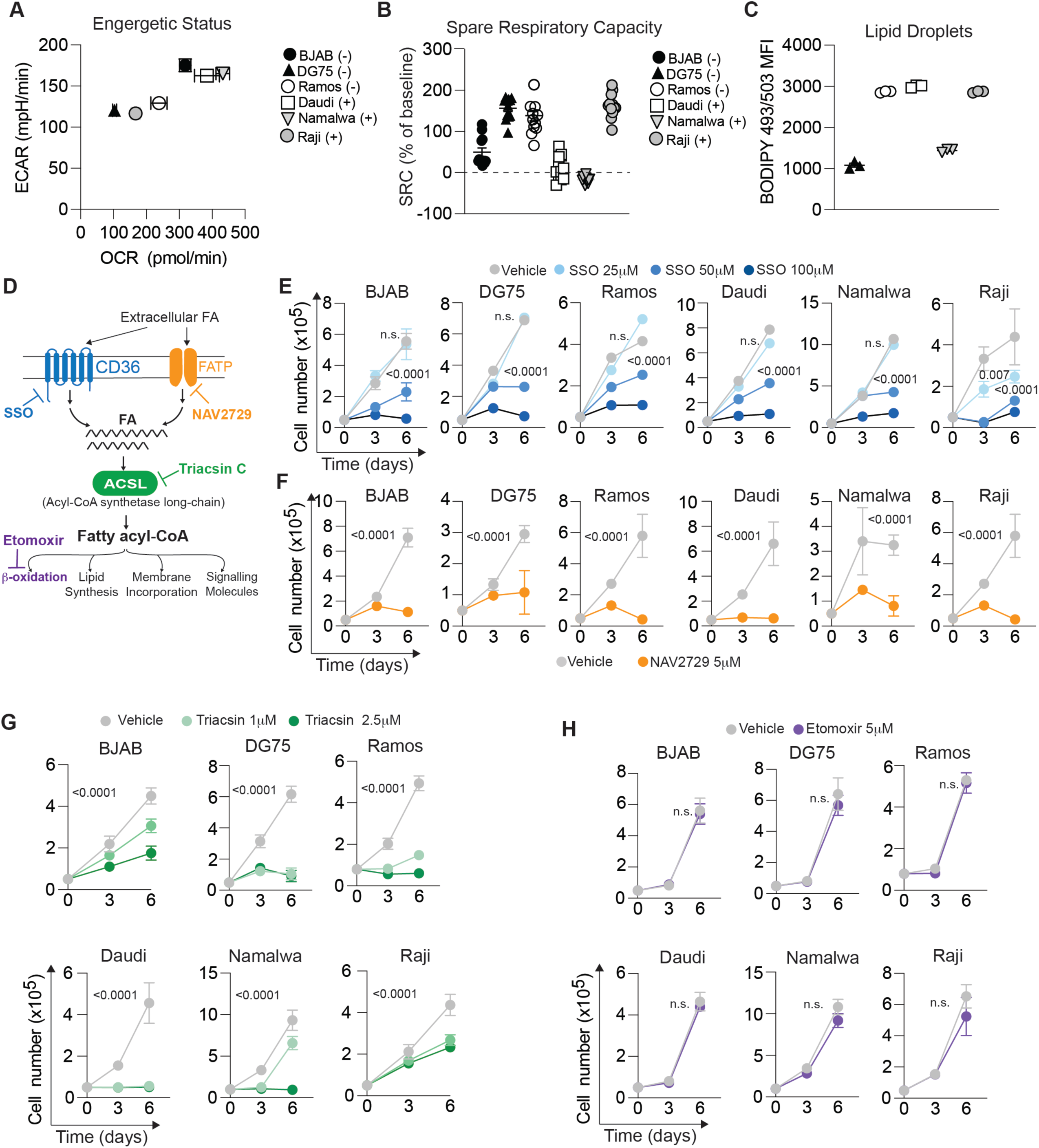
Burkitt lymphoma cells exhibit metabolic dependencies on fatty acid uptake and utilization. **(A–C)** Burkitt lymphoma cell lines historically associated with EBV positivity (Daudi, Namalwa, and Raji) or EBV negativity (BJAB, DG75, and Ramos) were evaluated for their metabolic and lipid profiles. **(A)** Energetic profiles determined by Seahorse Mito Stress Test and represented by oxygen consumption rate (OCR) versus extracellular acidification rate (ECAR). **(B)** Spare respiratory capacity, expressed as a percentage of basal respiration, determined following sequential treatment with oligomycin and FCCP during the Seahorse Mito Stress Test. **(C)** Neutral lipid content and lipid droplet accumulation assessed by BODIPY 493/503 staining and flow cytometry. (D–H) The requirement for fatty acid uptake and metabolism in Burkitt lymphoma cell proliferation was assessed using pharmacological inhibitors targeting distinct steps in lipid utilization. **(D)** Schematic depicting the lipid metabolic pathways and pharmacological inhibitors used, together with their respective targets. **(E)** Proliferation of Burkitt lymphoma cell lines in the presence of the CD36 inhibitor sulfo-N-succinimidyl oleate (SSO; 25, 50, and 100 µM). **(F)** Proliferation in the presence of the FATP2 inhibitor NAV-2729. **(G)** Proliferation in the presence of the long-chain acyl-CoA synthetase (ACSL) inhibitor Triacsin C. **(H)** Proliferation in the presence of the CPT1A inhibitor etomoxir. Data are presented as mean ± SD. Statistical significance was determined by two-way ANOVA followed by Tukey’s multiple-comparisons test.

Because EBV status did not predict major changes in mitochondrial or lipid metabolism, we next evaluated the essentiality of different nodes of fatty acid utilisation in Burkitt lymphoma (**Figure 1D**). In a dose-dependent manner, we found that inhibiting CD36, which imports saturated and unsaturated LCFA[33], with SSO[33] significantly reduced Burkitt lymphoma cell lines (**Figure 1E**), with an effect similar to inhibiting the fatty acid transporter FATP2 with NAV-2729 (Grassofermata[34]) (**Figure 1F**). Once saturated LCFAs enter the cell, they must first be activated by ACSLs before being directed toward downstream metabolic pathways, including fatty-acid oxidation or complex lipid synthesis[35]. Inhibition of this step with Triacsin C profoundly reduced proliferation, indicating the essential nature of lipid trafficking in Burkitt lymphoma (**Figure 1G**). In B-cell lymphoma cells, the entry and conversion of long-chain acyl-CoA into long-chain acylcarnitine in the mitochondria for FAO is facilitated by CPT1A and CPT1B[5]. Interestingly, inhibition of CPT1A with etomoxir[36] had no effect on B-cell lymphoma cell growth; however, inhibition of CPT1B with Oxfenicine[37] abrogated proliferation (**Figure 1H**, **Figure S1C**). Taken together, B-cell lymphomas exhibit a metabolic dependency on exogenous saturated fatty acids, as well as their downstream utilization for cell proliferation, including FAO via CPT1B.

### Sources of exogenous saturated fatty acids impact B-cell lymphoma proliferation

Saturated fatty acids – including MCFA and LCFA – can either come from the diet or from surrounding adipocytes in the tumour microenvironment of B-cell lymphoma. We first tested the effects of the MCFA Caprylic acid and the LCFA palmitic acid on B-cell lymphoma growth directly. The majority were sensitive to caprylic acid at higher concentrations, with Ramos and Daudi showing extreme sensitivity at lower doses (**Figure 2A**). However, all cell lines were sensitive to the highest dose of palmitic acid (100 µM), and most were sensitive even to the lower dose (**Figure 2B**). We next evaluated whether sensitivity to caprylic acid or palmitate correlated with changes in mitochondrial reactive oxygen species (mROS) using the probe MitoSOX. Palmitate, but not caprylic acid, altered mROS in a cell line-dependent manner, although these changes did not explain differential fatty acid sensitivity (**Figures S2A and B**). To further explore the impact of palmitate on mitochondrial function, we performed a Seahorse MitoStress test under low-glucose conditions with and without palmitate. While 4 of the 6 cell lines increased their basal OCR in response to palmitate, the least sensitive cell lines (Ramos and Daudi) were distinguished from the sensitive ones by a higher spare respiratory capacity, or mitochondrial flexibility (**Figure 2C and D**). Overall, these data indicate that B-cell lymphoma cells are sensitive to exogenous palmitate, possibly because they have a limited capacity to adapt to palmitate, possibly due to reduced mitochondrial fitness.

**Figure 2:**
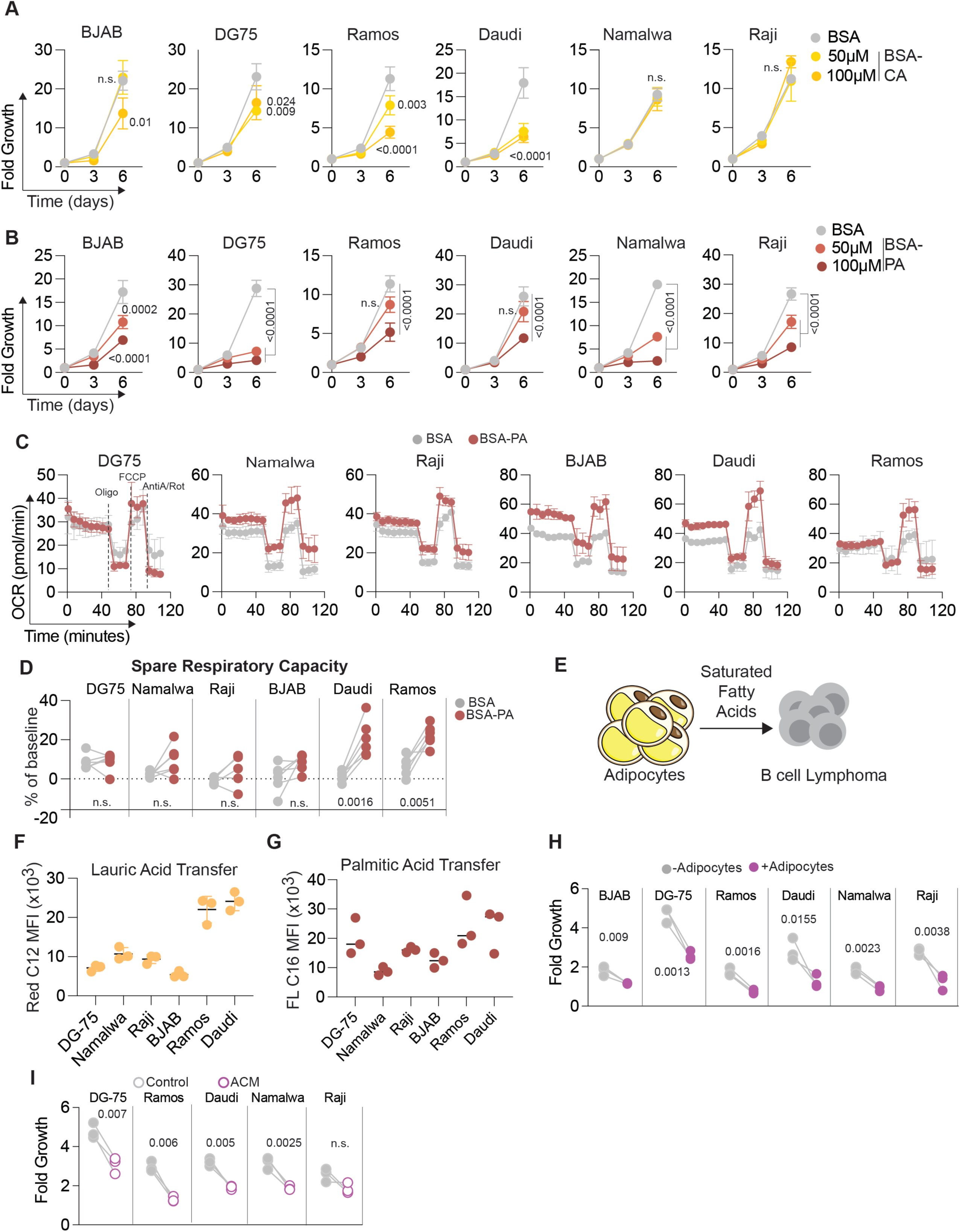
Exogenous saturated fatty acids and adipocytes crosstalk differentially affect B-cell lymphoma cell growth. **(A–B)** Proliferation of B-cell lymphoma cell lines cultured in the presence of increasing concentrations (50 and 100 μM) of the medium-chain fatty acid caprylic acid (CA; A) or the long-chain fatty acid palmitic acid (PA; B). **(C)** Mitochondrial function was assessed by Seahorse extracellular flux analysis using a Mito Stress Test under fatty acid oxidation conditions in the presence or absence of 50 μM PA. Oxygen consumption rate (OCR) was measured at baseline and following sequential injection of oligomycin, FCCP, and antimycin A/rotenone. **(D)** Spare respiratory capacity (SRC), expressed as a percentage of baseline respiration, calculated from the Mito Stress Test following oligomycin and FCCP treatment. **(E)** Schematic representation of the adipocyte–B-cell lymphoma co-culture system used to assess fatty acid transfer from adipocytes to lymphoma cells. **(F)** Transfer of the medium-chain fatty acid lauric acid from adipocytes to B-cell lymphoma cells, assessed using the fluorescent fatty acid analogue BODIPY™ 558/568 C12 (BODIPY Red C12). **(G)** Transfer of the long-chain fatty acid palmitic acid from adipocytes to B-cell lymphoma cells, assessed using fluorescent C16 (FL C16). **(H)** B-cell lymphoma proliferation, expressed as fold growth from day 0 to day 3, following culture in the presence or absence of adipocytes. **(I)** B-cell lymphoma proliferation, expressed as fold growth from day 0 to day 3, following culture in the presence or absence of adipocyte-conditioned medium (ACM). Data are presented as mean ± SD. For A–B, statistical significance was assessed by two-way ANOVA followed by Tukey’s multiple-comparisons test. Paired *t*-tests were used for the indicated paired comparisons in E, H, and I.

Adipocytes can serve as an exogenous source of fatty acids for solid cancers and leukemia[7; 38]. However, whether this occurs in B-cell lymphoma is unknown. Therefore, we measured whether adipocytes transfer MCFA and LCFA to B-cell lymphoma in co-culture systems. Using differentiated 3T3-L1s pre-stained with either the MCFA BODIPY (Red C12) or the LCFA BODIPY (FLC16), we measured their transfer to the six B-cell lymphoma cell lines (**Figure 2E**). Interestingly, the two MCFA-sensitive lines, Ramos and Daudi, showed the highest lauric acid uptake (**Figure 2F**). However, palmitate transfer from adipocytes was heterogeneous across cell lines (**Figure 2G**) and did not correlate with sensitivity. Patients with higher BMI in lymphoma have improved progression-free survival or overall survival[9; 39]. Because higher BMI is often associated with higher adiposity[40], we examined the direct effects of adipocytes on B-cell lymphoma proliferation. Strikingly, direct co-culture significantly impeded B-cell lymphoma growth (**Figure 2H**) and increased CD36 expression (**Figure S2C**), suggesting metabolic adaptation to the adipocyte-rich environment. These anti-proliferative effects were also recapitulated with adipocyte-conditioned media (**Figure 2I**), indicating that direct contact is not required. In a cell line-dependent manner, adipocyte-conditioned media can alter lipid uptake of the MCFA lauric acid and the LCFA palmitate, with ACM driving palmitate uptake more frequently (**Figure S2D and E**). Taken together, these data demonstrate that exogenous LCFA are a unique metabolic vulnerability in B-cell lymphoma and that lipid transfer from adipocytes impairs lymphoma cell growth.

### Dietary fat sources modulate B cell lymphoma growth, progression, and the tumour microenvironment

Diet manipulation in preclinical models, such as chronic obesogenic diets or fasting, has been shown to modulate B cell leukaemia growth and progression[13; 14; 41]. Paradoxically, however, higher body mass index (BMI) correlates with improved response to therapy in non-Hodgkin lymphoma and overall survival in diffuse large B-cell lymphoma (DLBCL)[9; 39]. We therefore turned to the orthotopic preclinical B-cell lymphoma model driven by Myc – the EμMyc model. We first confirmed that EμMyc B-cell lymphoma cells readily take up both MCFAs and LCFAs from the extracellular environment (**Figure 3A**). Given that adipocytes represent a physiologically relevant source of saturated fatty acids, we next examined lipid transfer from differentiated preadipocytes isolated from subcutaneous adipose tissue (SAT) and visceral adipose tissue (VAT). Both SAT- and VAT-derived adipocytes transferred MCFAs and LCFAs to EμMyc cells (**Figure 3B and C**). We next assessed whether these fatty acids differentially affected lymphoma cell growth. While caprylic acid (MCFA) had no detectable effect on EμMyc cell proliferation, exposure to palmitate (LCFA) significantly reduced proliferation (**Figure 3D and E**).

**Figure 3:**
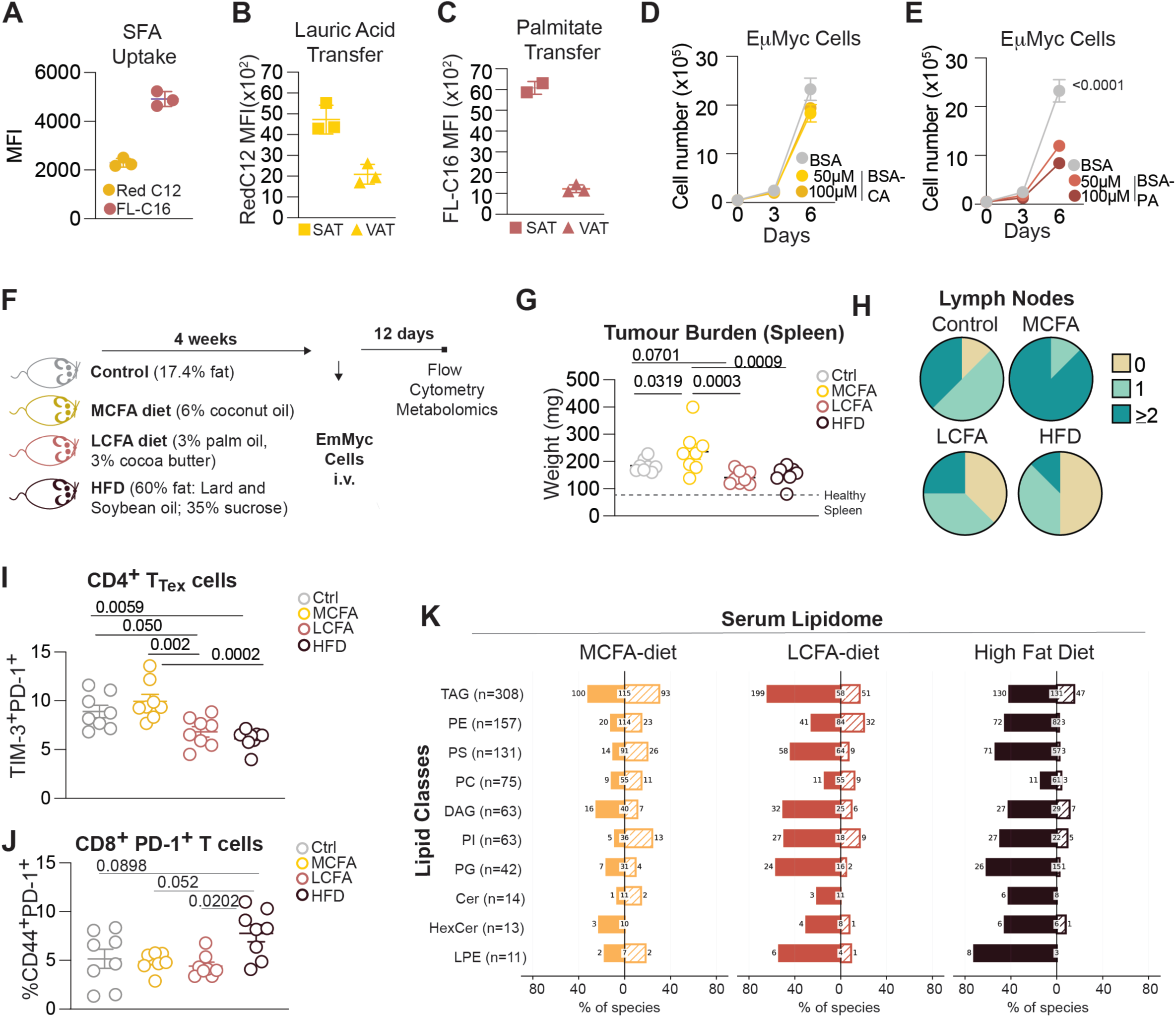
Dietary fatty acid composition regulates Eμ-Myc lymphoma growth, lipid metabolism, and T-cell exhaustion. **(A)** Uptake of medium- and long-chain fatty acids by Eμ-Myc lymphoma cells, assessed by flow cytometry using BODIPY Red C12 and FL C16, respectively. **(B-C)** Transfer of fatty acids from adipocytes differentiated from preadipocytes isolated from subcutaneous adipose tissue (SAT) or visceral adipose tissue (VAT) to Eμ-Myc lymphoma cells. Transfer of medium-chain BODIPY Red C12 **(B)** and long-chain FL C16 **(C)** fatty acids to Eμ-Myc cells was quantified by flow cytometry using the corresponding fluorescent fatty acid analogues. **(D-E)** Proliferation of Eμ-Myc lymphoma cells cultured in the presence of the medium-chain fatty acid caprylic acid (CA; D) or the long-chain fatty acid palmitic acid (PA; E). **(F)** Schematic representation of the orthotopic Eμ-Myc B-cell lymphoma model. Recipient male C57BL/6J mice were maintained on one of four diets: a control diet (17.4% fat); a medium-chain fatty acid (MCFA)-enriched diet, in which 6% of the control diet was replaced with coconut oil; a long-chain fatty acid (LCFA)-enriched diet, in which 3% of the control diet was replaced with cocoa butter and 3% with palm oil; or a high-fat obesogenic diet. **(G)**Lymphoma burden assessed by spleen weight at day 12 following Eμ-Myc lymphoma cell implantation (n=8 per condition). **(H)** Lymph node involvement assessed by lymph node size and number at endpoint (n=8 per condition). **(I)** Frequency of terminally exhausted CD4⁺ T cells, defined by co-expression of PD-1 and TIM-3, in spleens bearing Eμ-Myc lymphoma. **(J)** Exhaustion phenotype of splenic CD8⁺ T cells, assessed by PD-1 expression. **(K)** Serum lipidomic profiling by liquid chromatography-mass spectrometry (LC-MS) using the GIGA lipidomics platform. Lipid profiles were stratified according to major lipid classes, including triacylglycerols (TAG), phosphatidylethanolamines (PE), phosphatidylserines (PS), phosphatidylcholines (PC), diacylglycerols (DAG), phosphatidylinositols (PI), phosphatidylglycerols (PG), ceramides (Cer), hexosylceramides (HexCer), and lysophosphatidylethanolamines (LPE). For each experimental diet, lipid species are represented as changes relative to the control diet. Data are presented as mean ± SEM. Statistical significance was assessed by two-way ANOVA followed by Tukey’s multiple-comparisons test, as indicated.

We next tested whether dietary saturated fat composition impacts B cell lymphoma growth and progression *in vivo*. Recipient mice were placed in an isocaloric control diet, an MCFA-rich diet (6% of fat from coconut oil), and an LCFA-rich diet (3% from palm oil and 3% from cocoa butter), as well as an obesogenic diet for 4 weeks prior to EμMyc-cell transplantation (**Figure 3F**). At an intermediate time point (day 12), tumour burden, as assessed by splenic weight, was significantly increased in mice fed the MCFA-rich coconut oil diet, whereas mice fed the LCFA-rich palm oil/cocoa butter diet exhibited a modest reduction in tumour burden compared with control-fed mice (**Figure 3G**). However, the most striking finding was lymph node progression: a coconut oil-based diet accelerated lymph node progression, whereas LCFA-rich or high-fat diets limited it compared with controls (**Figure 3H**). Diet can also affect the tumour immune microenvironment (TIME), such as by modulating T cell function[11; 42]. Chronic consumption of a high-fat diet (HFD) has been associated with T-cell dysfunction and exhaustion[42]; however, whether shorter-term dietary exposures or changes in saturated fatty acid composition, independent of total dietary fat content, similarly influence T-cell exhaustion remains poorly understood. Therefore, we further examined T-cell exhaustion in the tumour microenvironment. Interestingly, both the LCFA diet and the HFD reduced terminal exhausted CD4 T cells compared to MCFA (**Figure 3I and Figure S3A**). However, exhausted PD-1+ CD8 T cells were modestly decreased in LCFA-fed mice compared with HFD, but did not differ from the other diets (**Figure 3J and Figure S3B**). Taken together, these data underpin a role for saturated fatty acid chain length in B cell lymphoma growth and in impacting the tumour immune microenvironment.

We next evaluated how saturated fat chain lengths change the full circulating lipidome compared with controls. An MCFA diet showed balanced increases and decreases across lipid species. However, both LCFA-rich and high-fat diets showed that there is a wide variety of decreases across a large range of lipid species: TAG, phosphatidylserine (PS), diacylglyceride (DAG), phosphatidylinositol (PI) and phosphatidylglycerol (PG) (**Figure 3K**). Species-level changes relative to control were compared between MCFA and HFD and between LCFA and HFD. This analysis revealed substantially greater concordance in both the magnitude and direction of lipid-species changes between LCFA and HFD than between MCFA and HFD, indicating that LCFA-rich feeding induces a circulating lipidomic profile that more closely resembles that of HFD (Figure S4A). LCFA-rich and obesogenic diets share a distinct circulating lipid signature that is largely absent following MCFA feeding and is characterized predominantly by depletion of TAG (**Figure S4B**). This shared response is particularly pronounced among PUFA-containing TAGs and PS, PI and PG species, indicating selective remodelling of circulating lipid composition rather than a uniform reduction in serum lipids (**Figure S4C**). The MCFA-rich diet – which exacerbated lymphoma growth and development – indicated a distinct circulating lipid profile compared with LCFA and high-fat diets, with a signature dominated by TAGs, PS, PC and PE species (**Figure S4D**). Deeper examination showed that the MCFA diet increased TAG species that were shorter and more saturated than in the other diets (**Figure S4E**). Collectively, dietary saturated fat chain length can profoundly remodel the lipidome and regulate lymphoma progression and the tumour microenvironment.

### ACSL1 as a biomarker of dietary response in B cell lymphoma

Precision nutrition as a co-therapy is a growing field in oncology[43]. Since a subset of B cell lymphoma cell lines were sensitive to the LCFA palmitate, we evaluated protein expression of metabolic regulators – c-MYC, Lipin-1, ACLY, LDHA, and ACSL1 – to identify a biomarker of response. Strikingly, the B cell lymphoma cell lines most sensitive to palmitate had lower ACSL1 expression than the more resistant lines (**Figure 4A**). Analysis of lymphoma patient cohorts using the Prediction of Clinical Outcomes from Genomic (PRECOG) database[44; 45] revealed that low ACSL1 expression was associated with reduced survival compared with high ACSL1 expression in DLBCL and, to a lesser degree, in Burkitt lymphoma (**Figure 4B and C**).

**Figure 4:**
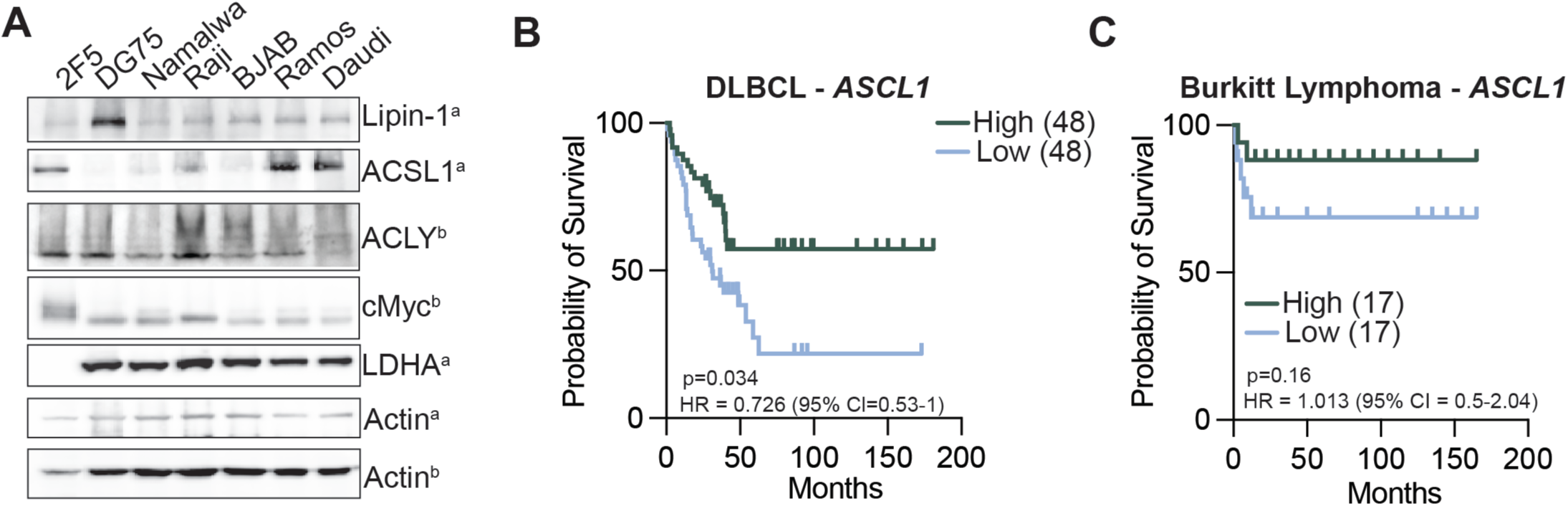
ACSL1 expression is associated with palmitate sensitivity and clinical outcome in B-cell lymphoma. **(A)** Protein lysates from murine Eμ-Myc lymphoma cells and human B-cell lymphoma cell lines (DG75, Namalwa, Raji, BJAB, Ramos, and Daudi) were analysed by immunoblotting for proteins involved in lipid metabolism and metabolic regulation, including lipin-1, ACSL1, ACLY, c-MYC, and LDHA. Human B-cell lymphoma cell lines are displayed in order of sensitivity to palmitic acid. β-actin was used as a loading control. Matching letters indicate proteins and corresponding loading controls derived from the same membrane. **(B-C)** Kaplan–Meier survival analysis of patients with diffuse large B-cell lymphoma (DLBCL; B) and Burkitt lymphoma **(C)**, stratified by low or high ACSL1 expression, using publicly available PRECOG gene expression and survival datasets.

Since *in vitro* and *in vivo* data indicated that ACSL1 low-expressing B-cell lymphomas are sensitised to LCFA, this could open a window of opportunity to provide a diet rich in LCFA to improve patient survival; we turned to mathematical modelling of this response.

### LCFA-enriched diets predicted to improve outcomes in Burkitt lymphoma with low ACSL1 expression

To quantify the effects of LCFAs on BL cell proliferation and predict the potential of an LCFA-enriched diet to improve outcomes in low-ACSL1-expressing lymphomas, we developed and calibrated a coupled Burkitt Lymphoma growth and inhibitor-response model (see Eqs. 1, 2, and 4 in the Methods, **Figure 5A**). Using our integrated model, we generated virtual individuals with either high- or low-ACSL1 expression Burkitt Lymphoma (BL) and predicted their tumour growth under a standard diet, and two diets with increased LCFA intake (Diets 1 and 2). Diets 1 and 2, which comprised LCFA concentrations peaking between 25 and 35 μM and 90 and 100 μM, respectively, represent moderate and substantial increases in dietary LCFA intake relative to a standard diet (see Methods and **Figure S7**).

**Figure 5:**
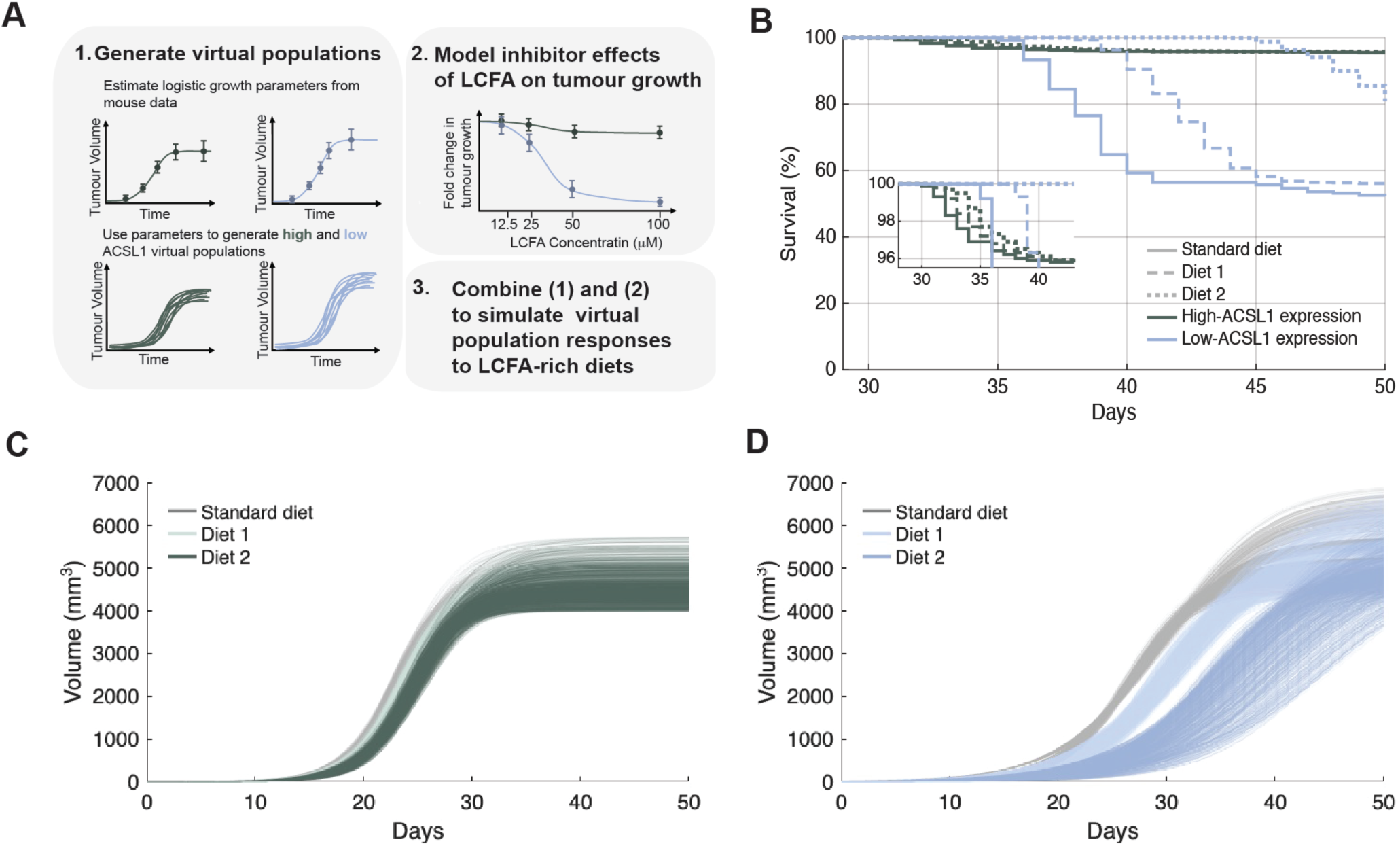
LCFA-enriched diets slow tumour growth and prolong survival in virtual Burkitt Lymphoma patients with low-ACSL1 expression. **(A)** Schematic overview of the methods used to generate the high-ACSL1 expression and low-ACSL1 expression virtual populations and predict their responses to LCFA-enriched diets. The workflow comprises three main steps: (1) generation of virtual populations, (2) modelling of the effects of LCFAs on tumour growth, and (3) integration of (1) and (2) to simulate virtual population responses to LCFA-enriched diets. **(B)** Simulated tumour growth trajectories for high-ACSL1 expression (left) and low-ACSL1 expression (right) virtual populations under a standard diet and two LCFA-enriched diets. Each population consists of 1,000 virtual patients, and each line represents the growth trajectory of an individual’s tumour under the indicated diet. **(C)** Kaplan-Meier survivorship curves for high- and low-ACSL1 expression virtual populations across the standard diet, Diet 1, and Diet 2. The inset magnifies the region where differences between diets in the high-ACSL1 population. **(B-C)** Standard diet: LCFA concentrations ≤5 μM. Diet 1: peak LCFA concentrations between 25 and 35 μM. Diet 2: peak LCFA concentrations between 90 and 100 μM (see Methods).

In the high-ACSL1 virtual population, both diets only slightly delayed tumour growth, resulting in minimal differences from a standard diet. However, our mathematical model predicted that LCFA-enriched diets could meaningfully slow tumour growth in the low-ACSL1 population (**Figure 5A**). We next related predicted tumour growth to survival probability by defining a lethality threshold of 5200 mm^3^ (see Methods). Although predicted survival probabilities did not precisely match the PRECOG data (**Figure 4C**), our simulations successfully captured the poorer overall survival in the low-ACSL1 group relative to the high-ACSL1 group. Indeed, while Diet 1 was predicted to increase survival modestly (from 52.3% to 55.3%), Diet 2 increased survival from 52.3% to 80.9%, a near 30 percentage-point improvement (**Figure 5C**). In contrast, LCFA enrichment had minimal effects on survival in the high-ACSL1 population, with model-predicted survival curves that were nearly indistinguishable, and endpoint survival rates of 95.5%, 95.6%, and 95.7% for a standard diet, Diet 1, and Diet 2, respectively (**Figure 5B**). Taken together, our model’s predictions suggest that increased dietary LCFA intake could slow BL progression and improve survival over a fixed period in patients with low-ACSL1 expression. This demonstrates the potential of an LCFA-rich diet as an adjunctive strategy to improve outcomes in this specific subgroup.

## Discussion

Dietary fat is generally considered in terms of its abundance or association with obesity; however, individual fatty acids have distinct metabolic properties that may differentially influence tumour cells and the surrounding immune microenvironment[11]. Here, we identify saturated fatty acid chain length as an important determinant of B-cell lymphoma metabolism and progression. B-cell lymphoma cells were dependent on pathways supporting exogenous fatty acid uptake and utilization, yet displayed strikingly different responses to medium- and long-chain saturated fatty acids. This distinction extended *in vivo*, where an MCFA-rich diet accelerated lymphoma progression, whereas an isocaloric LCFA-rich diet limited disease dissemination. Notably, LCFA-rich feeding generated changes in the circulating lipidome that closely resembled those induced by an obesogenic diet. Together, our findings suggest that the biological consequences of dietary fat cannot be explained solely by total fat intake and instead point to fatty acid chain length and tumour lipid-handling capacity as key determinants of lymphoma progression.

A central finding of this study is that B-cell lymphoma cells depend on exogenous lipid acquisition and processing. Pharmacological disruption of CD36- and FATP-mediated fatty acid uptake, as well as ACSL-dependent fatty acid activation, markedly impaired lymphoma cell proliferation. Surprisingly, CPT1A inhibition had little effect, whereas targeting CPT1B strongly suppressed proliferation. The differential requirement for CPT1 isoforms is particularly intriguing given the predominant focus on CPT1A in cancer metabolism. However, in acute myeloid leukaemia cells, CPT1B is overexpressed and drives FAO-mediated metabolic reprogramming[46]. The key difference is that CPT1A has a higher affinity for carnitine than CPT1B, but CPT1B is inhibited by malonyl-CoA (product of fatty acid synthesis) at lower concentrations[47]. Therefore, lymphoma cells may rely more on CPT1B because it is more tightly coordinated with metabolic needs for fatty acid synthesis.

Our findings reveal a context-dependent relationship between adipocytes and lymphoma cells in which adipocyte-derived factors reduce B cell lymphoma growth. This is the opposite of many other cancer types, such as pancreatic and ovarian cancer, where adipocytes provide a tumour-supportive network[6; 8]. Paradoxically, in B cell lymphoma, higher BMI have been associated with improved responses and overall survival in obese patients[9; 39]. Our data could provide a new metabolic framework in which increased saturated lipid metabolism induces metabolic weakness in B cell lymphoma and could be exploited via dietary intervention. We also found that ACSL1 expression inversely correlated with LCFA sensitivity. While ACSL1 is essential to B cell lymphoma, its expression could be used as a biomarker of response to exogenous LCFA. The orthotopic EμMyc model, which has moderate ACSL1 expression, is sensitive to diets rich in saturated LCFA, similar to *in vitro* responses. Using mathematical modelling to test the sensitivity of diets richer in LCFA and ACSL1 expression stratified as high and low, we found, as proof of principle, that we have indeed identified a metabolic vulnerability of ACSL-low-expressing lymphomas to dietary saturated LCFA. However, this would need further exploration in preclinical models.

Exogenous lipids could exert their effects on B-cell lymphomas through various mechanisms. One obvious mechanism could be changes in membrane composition that affect membrane fluidity. Serum lipidomics suggest that changes in lipid partitioning induce a converging phenotype in the LCFA-rich and obesogenic diet, where 18:0-containing phospholipids are depleted, suggesting they are diverted away from structural membrane lipids. Membrane fluidity and order are biophysical properties of the plasma membrane determined in part by lipid composition, with changes in lipid packing influencing membrane organisation, receptor signalling, and cellular function[48; 49]. Membrane fluidity is essential for BCR-dependent signal transduction in B cells, and modifying membrane order in B cell lymphoma regulates cell death[50]. The tumour microenvironment can also shape membrane fluidity in lymphoma cells, with higher membrane order found in the spleen than in ascites[51]. Identifying novel approaches to modify membrane order could improve responses to therapies, since changes in membrane fluidity are often associated with therapy resistance[52; 53]. Taken together, our data suggest that dietary intervention for B-cell lymphomas could be used in the context of precision nutrition and potentially co-administered to potentiate responses to existing therapies.

In conclusion, our study reveals that saturated fatty acids are not metabolically interchangeable in B-cell lymphoma. Instead, fatty acid chain length determines how dietary and adipocyte-derived lipids interact with tumour metabolic capacity, circulating lipid composition and anti-tumour immunity. LCFA exposure creates a metabolic vulnerability in a subset of lymphoma cells, whereas MCFA-rich feeding promotes lymphoma progression and generates a distinct circulating lipid environment. By identifying ACSL1 as a potential determinant of LCFA adaptation, these findings establish a mechanistic framework linking dietary lipid composition to tumour metabolism and suggest that exploiting lipid-handling vulnerabilities may provide a foundation for metabolically informed precision nutrition in B-cell lymphoma. More broadly, our findings argue against considering saturated dietary fats as a homogeneous nutritional class and suggest that defining the molecular composition of dietary fat, together with tumour-specific metabolic capacity, may provide a framework for developing precision nutritional strategies in B-cell lymphoma.

## Acknowledgements

We acknowledge the Animal Facility and Flow Cytometry Facility at the Centre de recherche de l’Hôpital Maisonneuve-Rosemont (CR-HMR) for their technical expertise and support. We are grateful to Drs. Jerry Pelletier and Francis Robert at McGill University for providing the Eμ-Myc cell lines and to the laboratory of Dr Frédérick A. Mallette for providing mouse embryonic fibroblasts. This work was supported by the Cancer Research Society Operating Grant (Award ID 1057230), the Université de Montréal Appui aux initiatives intersectorielles program, and the Cole Foundation. J.B. is supported by a Fonds de recherche du Québec (FRQ) Junior 1 salary award; E.B.G. by the Stavros Niarchos Foundation; B.N. by an FRQ Master’s studentship; and L.B. by a Cole Foundation postdoctoral fellowship.

## Author Contributions

J.B., F.Z, and M.C conceived and designed the study, contributed to experimental work and data interpretation, and J.B wrote the manuscript. A.L.C., B.N, and C.G performed experiments and analyzed data. M.P. and M.C. contributed to experimental design, and M.P. performed the mathematical modelling and wrote the corresponding sections of the manuscript. L.B. and A.L.C. performed the in vivo orthotopic experiments with assistance from M.L. and E.B.G. E.B.G. and I.G performed flow cytometry experiments. A.M provided expertise in culturing B-cell lymphoma cell lines and the EμMyc model. E.C.C provided samples, helped generate cell lines and provided input on the EuMyc model. B.N. contributed to manuscript preparation. All authors reviewed and approved the final version of the manuscript and agreed to its submission.

---

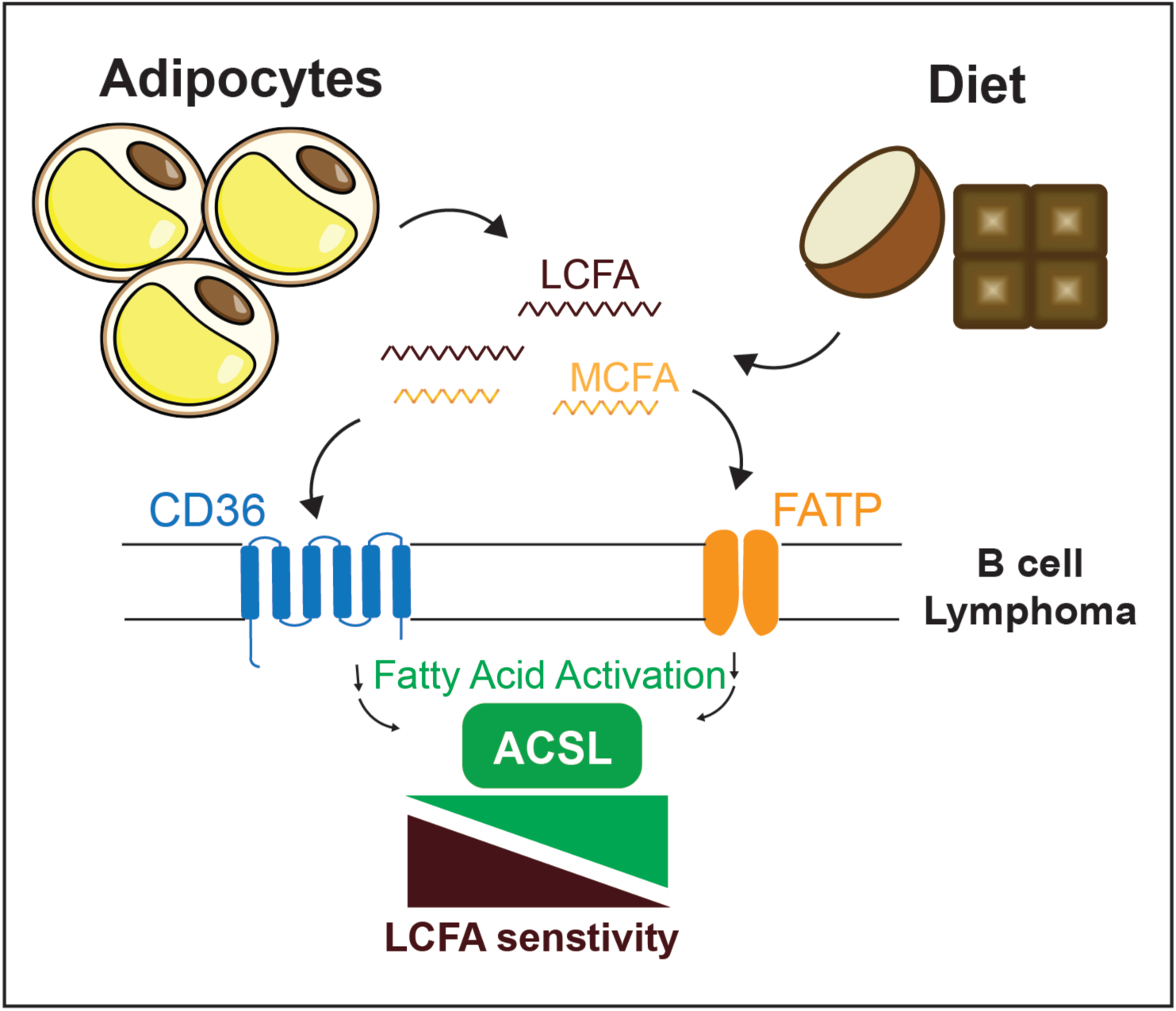

