## supplemental file for "Saturated fatty acid chain length shapes B cell lymphoma metabolism and progression"

**Supplementary Figures – Collin *et al.***

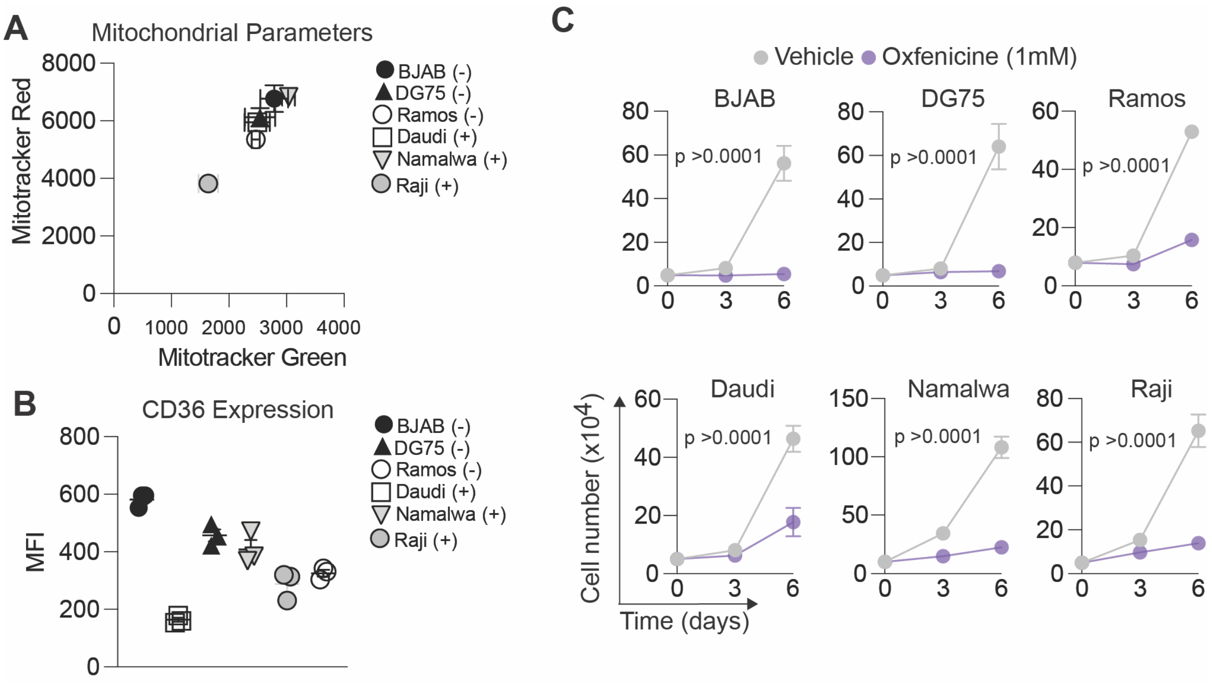


**Figure S1: Mitochondrial and lipid metabolic characteristics of B-cell lymphoma cell lines.** **(A)** Mitochondrial membrane potential and mitochondrial mass of B-cell lymphoma cell lines assessed by MitoTracker Red and MitoTracker Green staining, respectively, and flow cytometry. **(B)** CD36 expression across B-cell lymphoma cell lines measured by flow cytometry. **(C)** Proliferation of B-cell lymphoma cell lines in the presence of increasing concentrations of oxfenicine, an inhibitor of CPT1B-mediated fatty acid oxidation. Data are presented as mean ± SD. For **(C)**, statistical significance was determined by two-way ANOVA followed by Tukey's multiple-comparisons test.

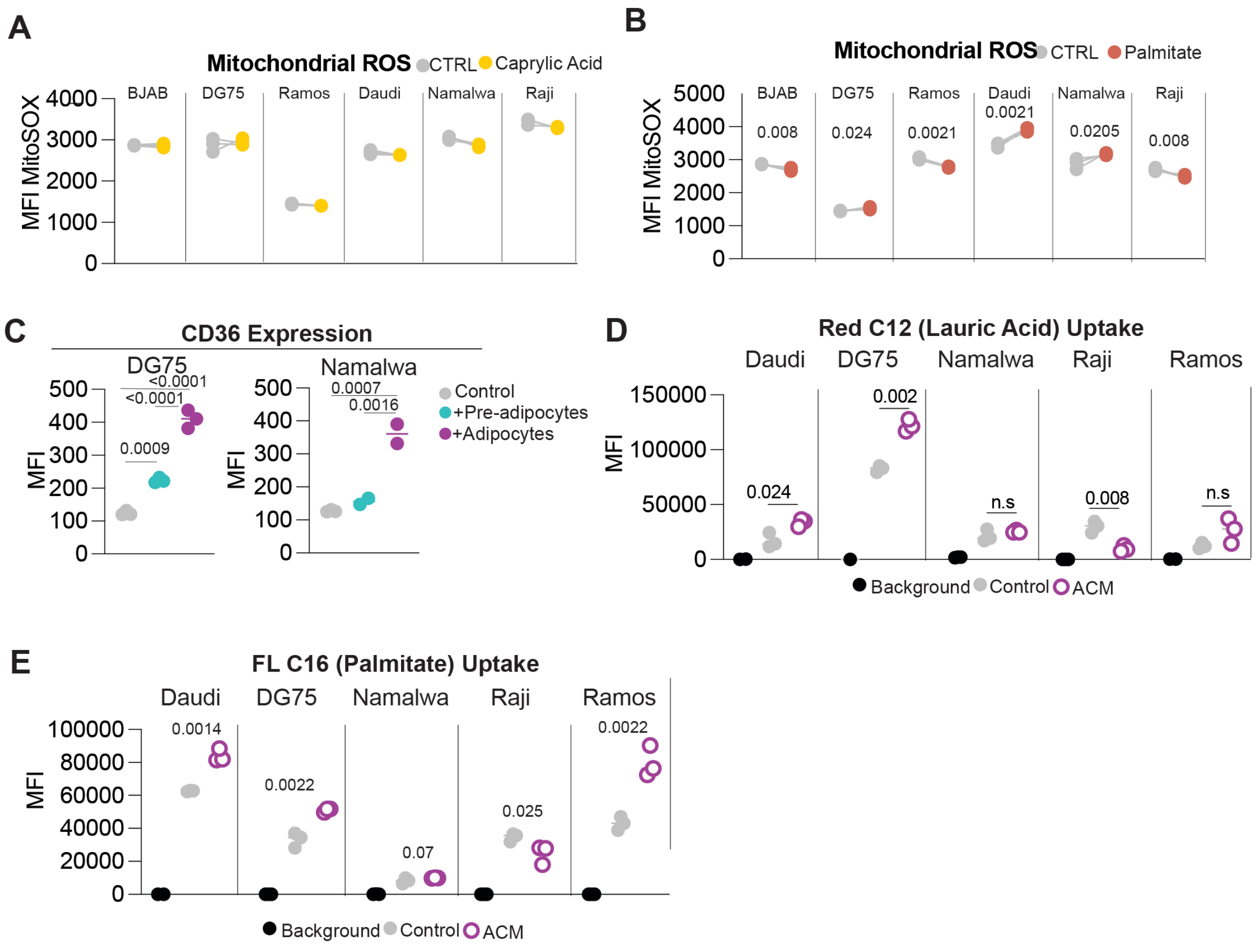


**Figure S2: Exogenous fatty acids and adipocyte-derived factors modulate lipid uptake and metabolism in B-cell lymphoma cells.**

**(A–B)** Mitochondrial reactive oxygen species (mROS) in B-cell lymphoma cell lines following 48 h exposure to fatty acids, assessed by MitoSOX staining and flow cytometry. **(A)** mROS production following exposure to caprylic acid (C8). **(B)** mROS production following exposure to palmitic acid (C16). **(C)** CD36 expression in B-cell lymphoma cells cultured alone or co-cultured with undifferentiated 3T3-L1 preadipocytes or differentiated 3T3-L1 adipocytes for 3 days, assessed by flow cytometry. **(D)** Lauric acid uptake by B-cell lymphoma cells cultured for 3 days under control conditions or with adipocyte-conditioned medium (ACM), assessed using BODIPY Red C12 and flow cytometry. **(E)** Palmitic acid uptake by B-cell lymphoma cells cultured for 3 days under control conditions or with ACM, assessed using FLC16 and flow cytometry. Data are presented as mean ± SD. Statistical significance was determined using paired two-tailed Student's *t*-tests.

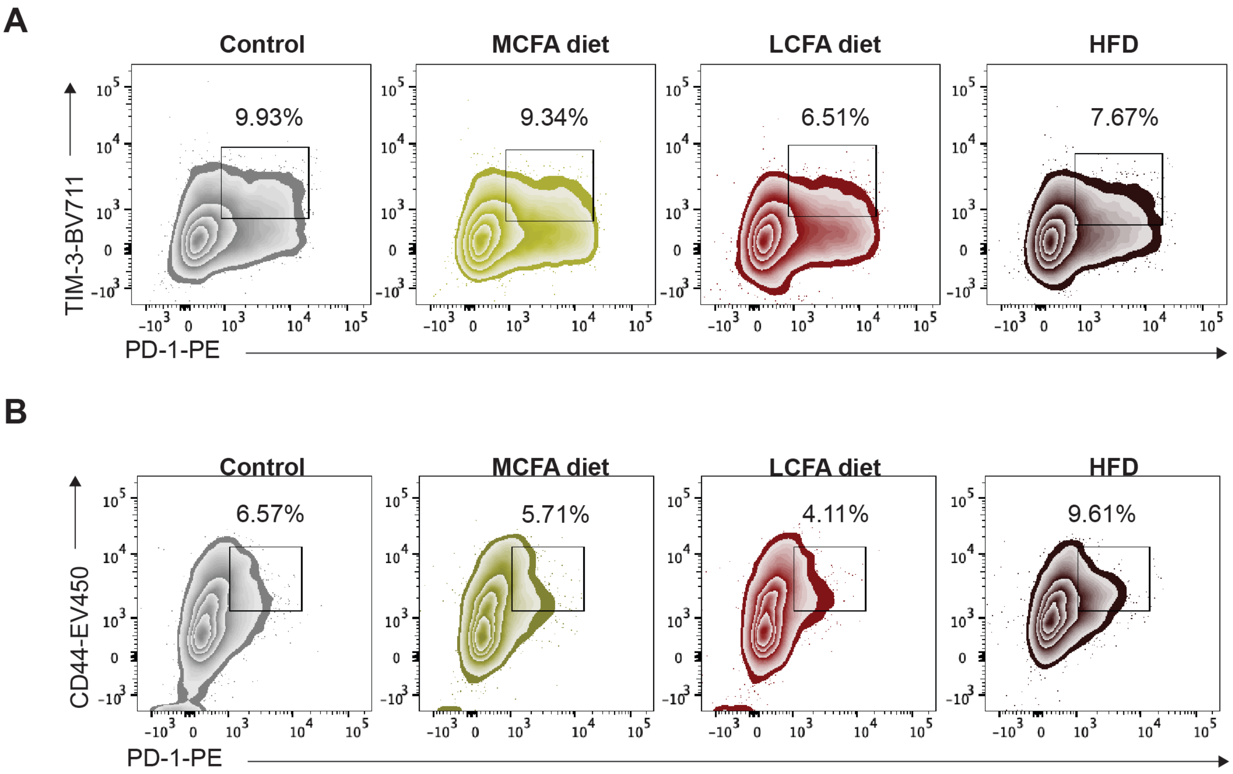


**Figure S3: Dietary saturated fat composition alters exhausted T-cell populations in splenic B-cell lymphoma.**

(**A-B**) Representative flow cytometry plots of exhausted T-cell populations within splenic tumours from mice maintained on a control diet, an isocaloric medium-chain fatty acid (MCFA)-rich diet, an isocaloric long-chain fatty acid (LCFA)-rich diet, or an obesogenic high-fat diet (HFD). **(A)** Representative flow cytometry plots showing terminally exhausted CD4 T cells identified by co-expression of PD-1 and TIM-3 within splenic tumours. **(B)** Representative flow cytometry plots showing exhausted CD8 T cells identified based on PD-1 expression within splenic tumours. Numbers indicate the percentage of cells within the indicated gates.

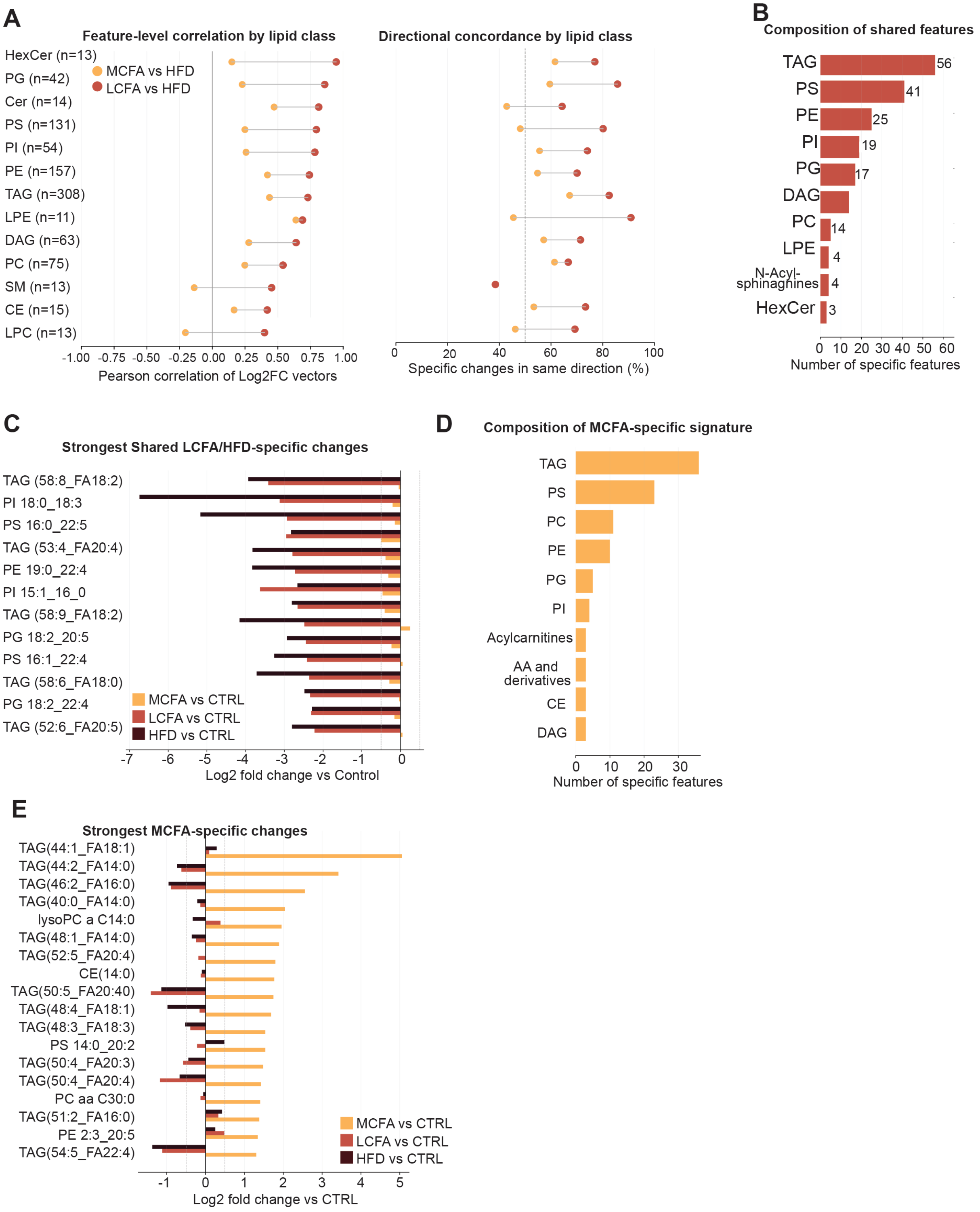


**Figure S4. LCFA-rich and high-fat diets induce shared circulating lipidomic changes that are distinct from the MCFA-rich diet.**

**(A)** Lipid class-specific similarity between dietary responses. Left, feature-level Pearson correlations of lipid-species log2 fold changes relative to control for MCFA versus HFD and LCFA versus HFD within each indicated lipid class. Right, directional concordance, defined as the percentage of lipid species within each class exhibiting changes in the same direction relative to control. The number of lipid species included in each class is indicated. **(B)** Lipid-class composition of features exhibiting shared changes between the LCFA-rich diet and HFD, showing the number of shared lipid species within each class. **(C)** Lipid species exhibiting the greatest shared LCFA/HFD-specific changes. Bars represent log2 fold changes relative to control in mice maintained on MCFA-rich, LCFA-rich, or high-fat diets. **(D)** Lipid-class composition of the MCFA-specific circulating lipidomic signature, represented as the number of MCFA-specific features belonging to each lipid class. **(E)** Lipid species exhibiting the strongest MCFA-specific changes. Bars represent log2 fold changes relative to control for the MCFA-rich, LCFA-rich, and high-fat diet groups. TAG, triacylglyceride; PS, phosphatidylserine; PE, phosphatidylethanolamine; PI, phosphatidylinositol; PG, phosphatidylglycerol; DAG, diacylglyceride; PC, phosphatidylcholine; LPE, lysophosphatidylethanolamine; HexCer, hexosylceramide; SM, sphingomyelin; CE, cholesteryl ester; LPC, lysophosphatidylcholine.

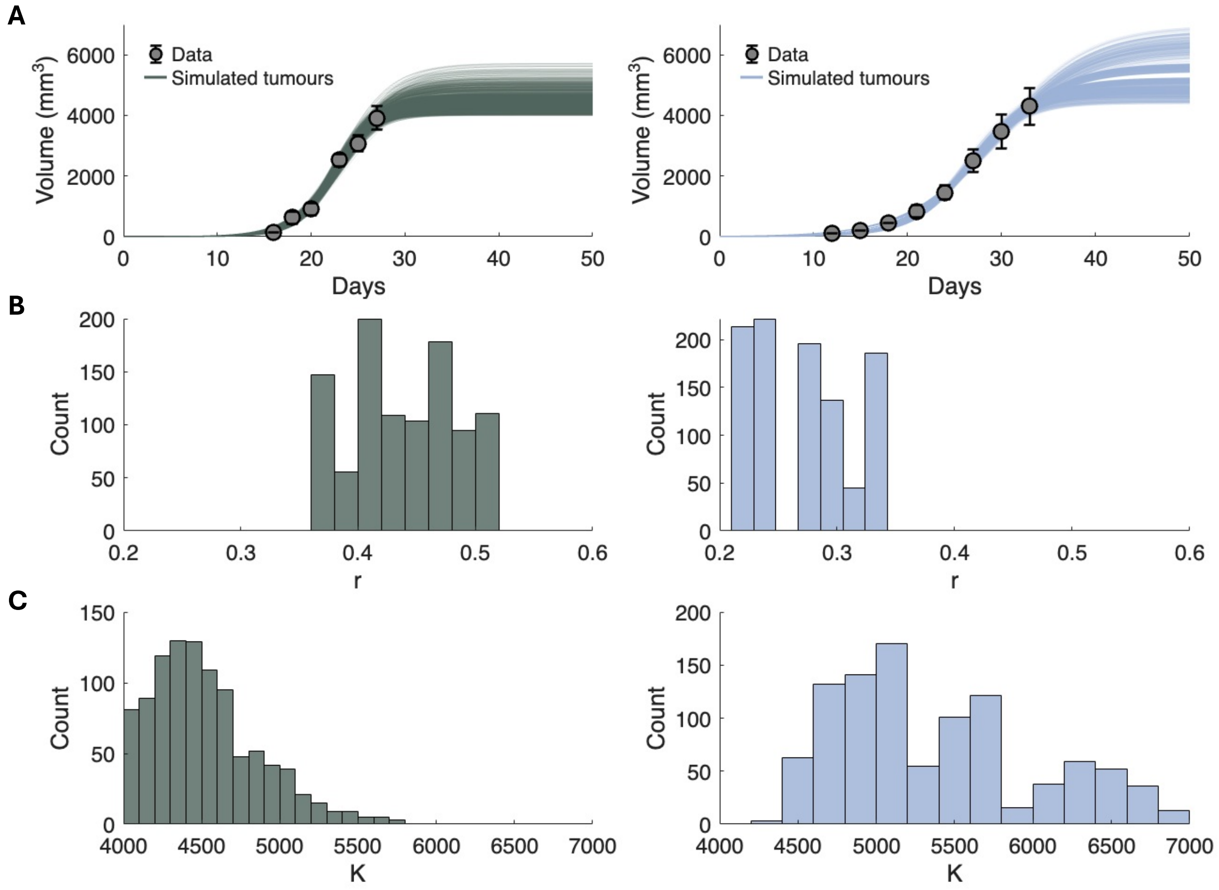


**Figure S5: Virtual Burkitt lymphoma populations recapitulate mouse tumour-growth data and capture inter-individual heterogeneity. (A)** Simulated tumour growth (Eq. 4 in the Main Text) trajectories of 1,000 high-ACSL1 expression (left) and 1,000 low-ACSL1 expression (right) virtual mice under untreated conditions. Grey dots represent the tumour growth data used to generate the virtual populations[1,2] (see Methods in the Main Text). That the bulk of simulated trajectories pass through the standard deviation bars indicates good agreement between model predictions and the data. Distributions of the **(B)** growth rate, $r$, and **(C)** maximum tumour volume, $K$, in the high- (left) and low- (right) ACSL1-expression virtual populations.

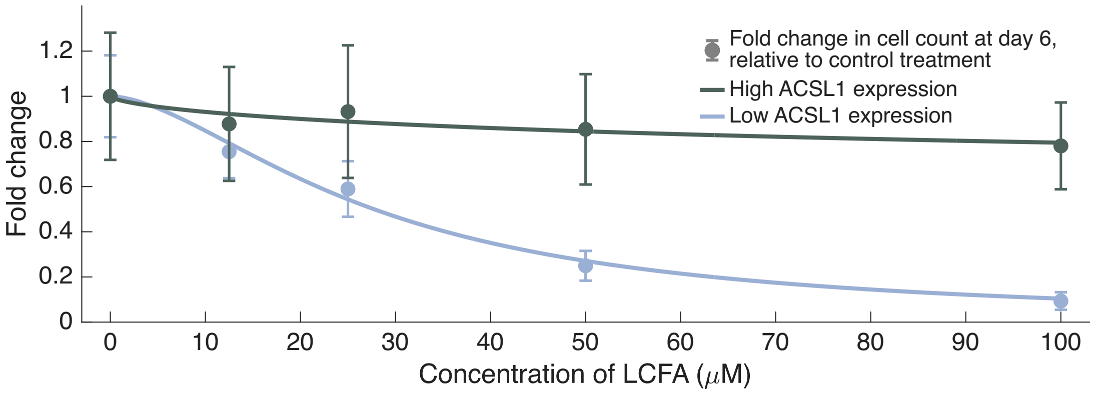


**Figure S6: Low-ACSL1 expression Burkitt lymphomas show high sensitivity to LCFAs.** The classic inhibitor-response curve (Eq. 2 in the Main Text) was used to model the effect of LCFAs on proliferation in Burkitt lymphoma cell lines, with LCFAs treated as the inhibitor and response measured as fold change in cell count relative to the untreated control (see Methods in the Main Text). Data points represent fold change values for each treatment condition, grouped by ACSL1 expression of the cell lines: high-expression cells (n = 6, green) and low-expression cells (n = 6, blue; see Figure 2B in the Main Text). Solid lines represent the best-fit inhibitor-response curve for each expression group. Fitted parameter values for both these curves are presented in Table S1.

| | $E_{min}$ | $E_{max}$ | $IC50$ | $h$ |
|---|---|---|---|---|
| **Low ACSL1 expression** | 1 (fixed) | $1.7 \cdot 10^{-12}$ | 28 | 1.7 |
| **High ACSL1 expression** | 1 (fixed) | 0.53 | 150 | 0.65 |

**Table S1: Estimated parameter values of inhibitor-response model.** See Eq. 2 and the Methods in the Main Text for description of parameters and method of parameter estimation.

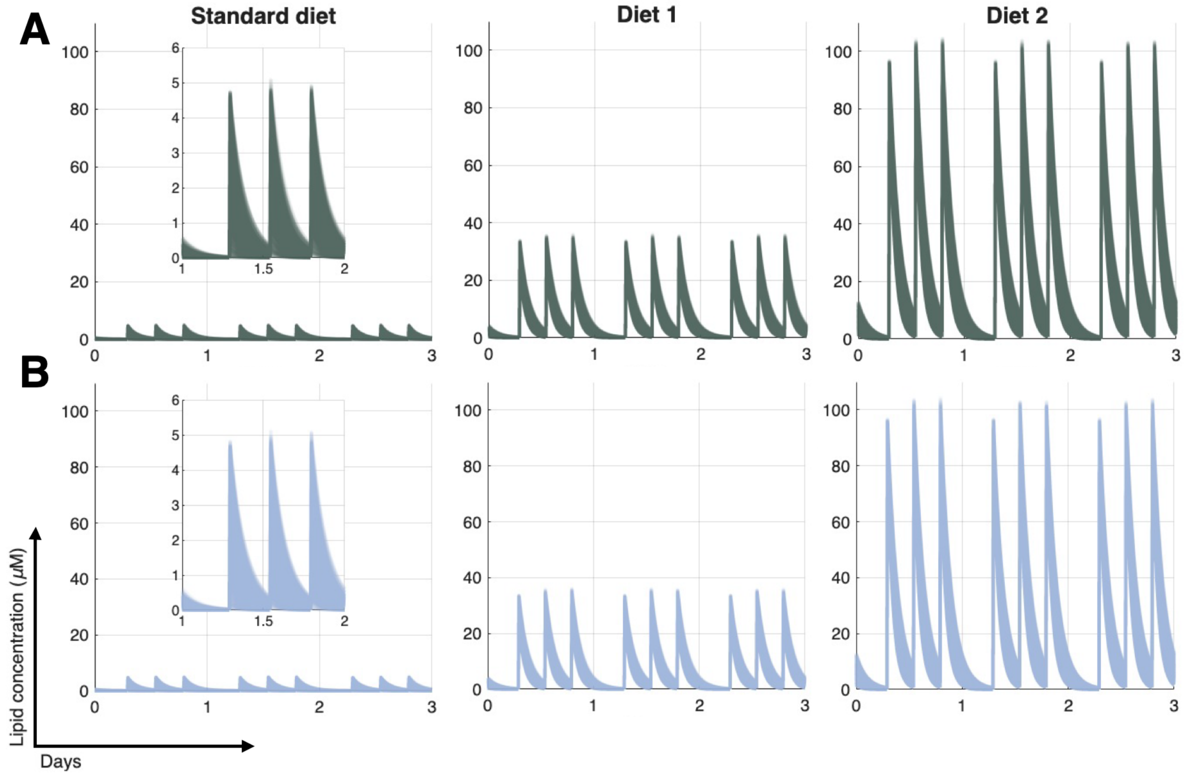


**Figure S7: Simulated diets have time-varying LCFA concentrations.** LCFA concentrations over time following the three diets studied in the high- **(A)** and low- **(B)** ACSL1 virtual populations. In each, LCFA concentrations rise instantaneously following each of three simulated daily meals (evenly spaced over a 12-hour feeding window followed by a 12-hour fasting period) then decay exponentially between meals. Each column corresponds to one of the studied three diets. **Left to right:** Standard diet (low, baseline intake of LCFAs), Diet 1 (moderate increase in LCFA intake), and Diet 2 (substantial increase in LCFA intake).

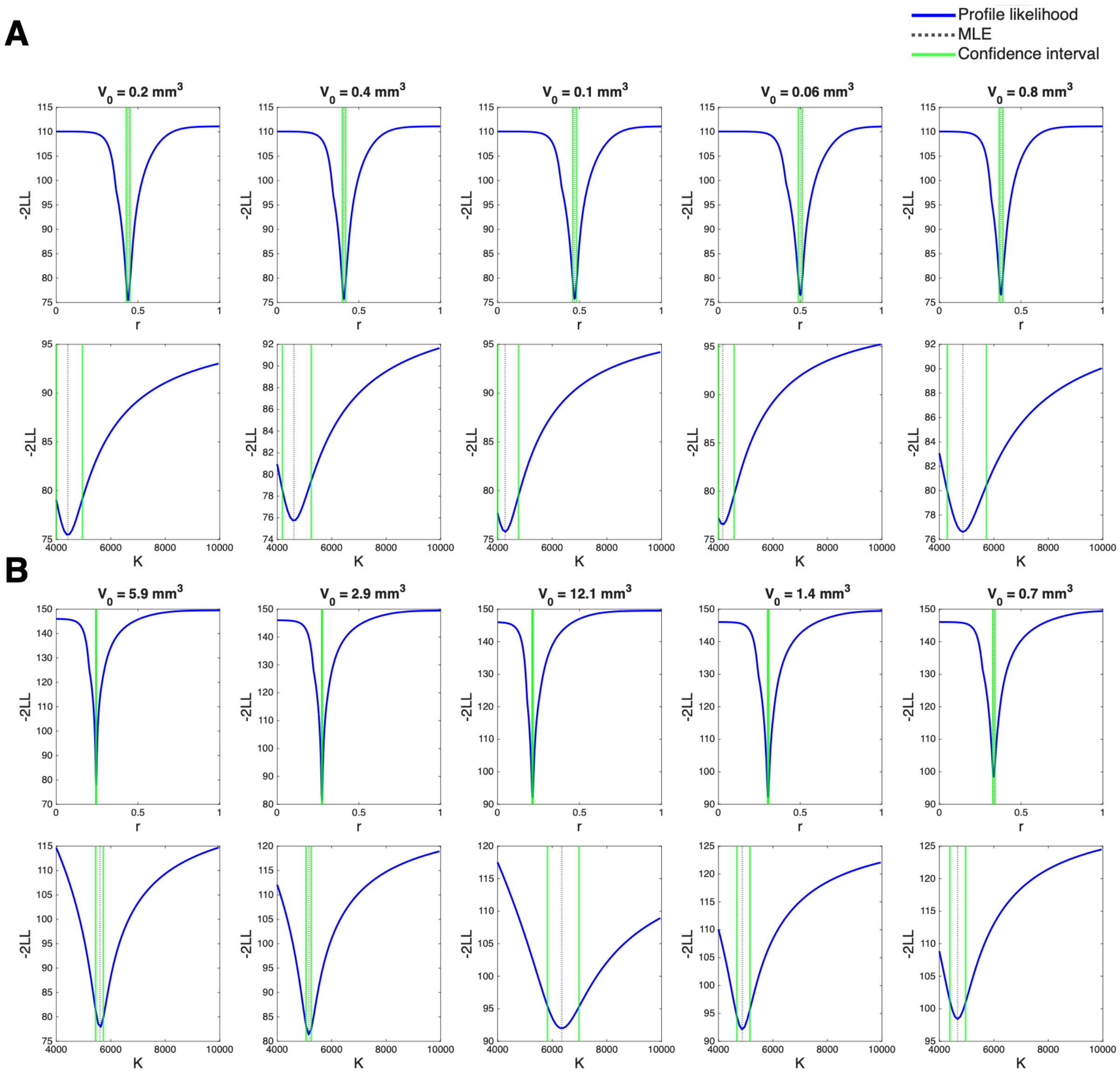


**Figure S8: Profile likelihoods for high- and low-ACSL1 expression groups.** Profile likelihoods for the high- **(A)** and low- **(B)** ACSL1 expression data. **(A-B)** Top row: PLs for the intrinsic growth rate, $r$. Bottom row: PLs for maximal tumour volume, $K$. All profiles display finite confidence intervals and clear minimums, demonstrating practical identifiability. The profile likelihood method[3] was employed to verify practical identifiability and estimate 95% confidence intervals for the parameter values in the logistic growth model (Eq. 1 in the Main Text). The results from this analysis were used to generate the virtual populations.

| | $V_0$ | $r$ | | $K$ | |
|---|---|---|---|---|---|
| | | Lower bound | Upper bound | Lower bound | Upper bound |
| High ACSL1 expression | 0.2 | 0.4284 | 0.4505 | 4000 | 4961 |
| | 0.4 | 0.3984 | 0.4184 | 4192 | 5249 |
| | 0.1 | 0.4585 | 0.4825 | 4000 | 4769 |
| | 0.06 | 0.4885 | 0.5125 | 4000 | 4577 |
| | 0.8 | 0.3664 | 0.3884 | 4288 | 5730 |
| Low ACSL1 expression | 5.9 | 0.2422 | 0.2442 | 5441 | 5730 |
| | 2.9 | 0.2723 | 0.2763 | 5057 | 5249 |
| | 12.1 | 0.2102 | 0.2162 | 5826 | 6979 |
| | 1.4 | 0.3003 | 0.3083 | 4673 | 5153 |
| | 0.7 | 0.3263 | 0.3403 | 4384 | 4961 |

**Table S2: Initial conditions and corresponding parameter 95% confidence intervals for each expression group.** Initial conditions and confidence intervals were later used to generate our virtual populations (see Methods in the Main Text)
